# On the Comparison of LGT networks and Tree-based Networks

**DOI:** 10.1101/2025.11.20.689557

**Authors:** Bertrand Marchand, Nadia Tahiri, Olivier Tremblay-Savard, Manuel Lafond

## Abstract

Phylogenetic networks are widespread representations of evolutionary histories for taxa that undergo hybridization or Lateral-Gene Transfer (LGT) events. There are now many tools to reconstruct such networks, but no clearly established metric to compare them. Such metrics are needed, for example, to evaluate predictions against a simulated ground truth. Despite years of effort in developing metrics, known dissimilarity measures either do not distinguish all pairs of different networks, or are extremely difficult to compute. Since it appears challenging, if not impossible, to create the ideal metric for all classes of networks, it may be relevant to design them for specialized applications. In this article, we introduce a metric on LGT networks, which consist of trees with additional arcs that represent lateral gene transfer events. Our metric is based on edit operations, namely the addition/removal of transfer arcs, and the contraction/expansion of arcs of the base tree, allowing it to connect the space of all LGT networks. We show that it is linear-time computable if the order of transfers along a branch is unconstrained but NP-hard otherwise, in which case we provide a fixed-parameter tractable (FPT) algorithm in the level. We implemented our algorithms and demonstrate their applicability on three numerical experiments.

**Full online version:** https://www.biorxiv.org/content/10.1101/2025.11.20.689557

## 1 Introduction

Phylogenetic networks are often seen as more accurate representations of evolution than trees, especially for species that undergo hybridization or lateral gene transfer, and one can even argue that the “tree of life” hypothesis should be updated to incorporate network-like structures [1]. Several approaches now aim to reconstruct networks, for example NeighborNet [2], PhyloNet [3], SNaQ [4], PhyNest [5], T-Rex [6], and more. Despite the importance of predicting such networks, there is still no clearly established method to evaluate the quality of the reconstructions. For phylogenetic *trees*, the Robinson-Foulds (RF) metric and extensions [7,8] are widely used, and despite some shortcomings are accepted as a good measure of similarity. Therefore, when a new tree prediction tool is developed, it can be evaluated against a gold standard or simulated dataset using RF, providing insights on its strengths and weaknesses.

In the case of networks, this is not so easy. Several network measures have been attempted in the last two decades, but all of them have drawbacks. This includes structural measures that compare small components of the networks, such as softwired or hardwired clusters [9,10,11], rooted triples and trinets [12,13,14], displayed trees [15], and others [9,16], as well as metrics based on counting paths, such as the *µ*-distance [17,18], which was recently generalized to semi-directed networks [19]. All of these metrics suffer from the fact that, in all generality, they may say two networks are the same when they are actually different (see e.g. [9,16]). Another family of measures are operational metrics, which are based on a number of operations needed to transform one network into the other (e.g., SPR [20,21,22], NNI [23,24], contractions [25], or cherry-picking [26,27] operations). However, these tend to be very hard to compute and, to our knowledge, not yet used frequently in practice. Overall, it appears extremely difficult to design a “one-size-fits-all” metric for networks, and it may be more relevant to design metrics that are tailored for specific applications.

Towards this goal, our aim in this paper is to compare networks in which reticulations represent lateral gene transfer (LGT) events. Specialized approaches can be used to reconstruct such networks, since the problem can be formulated in terms of starting with a species *tree*, and inserting arcs between branches to represent genetic exchanges between co-existing species. This results in what is called an *LGT network*, which consisting of the known original *base tree* and extra *transfer arcs* [28]. There are many LGT network reconstruction approaches: they can be built using tri-nets [29]; using so-called *Duplication-Transfer-Loss (DTL) reconciliation* [30,31]; using orthology and xenology data [32]; or using the presence/absence of character traits to predict clades that exchanged characters [33,34]. Moreover, LGT networks are a specialized form of *tree-based networks*, which also consist of base trees with additional arcs, but in which the base tree is unknown [35]. As tree-based networks form a popular class that generalizes many others [36], there is also a need to develop metrics for them.

Even in the case of LGT networks, there appears to be no ideal comparison metric. As a result, LGT network predictors have mostly been evaluated in an ad hoc manner in previous work. For example in [34], LGT networks were reconstructed to predict bacterial transfers, notably interphylum transfers between Proteobacteria and Actinobacteria, and the results were compared to published studies on this set of species. Unfortunately, this methodology is qualitative and does not easily generalize to other datasets. As already mentioned, another way of reconstructing LGT networks is by computing *DTL reconciliations* between species and gene trees [31,37]. These infer events on gene trees and the predicted gene transfers can be used to add LGT arcs on the species tree. These approaches typically measure the accuracy of the events on the gene trees, but not the accuracy of the underlying species LGT network — which is more relevant in some contexts such as transfer highway identification.

### Our results

In this work, we address the aforementioned gaps by proposing a novel metric to compare LGT networks. We exploit the distinction between the base tree and the transfer arcs and compare these two components in an independent fashion. The “base tree components” of the networks are compared using the established Robinson-Foulds metric, which reduces the problem to comparing only the “transfer components”. An intuitive way of comparing the sets of transfer arcs in two networks would be to count the number of “same” transfers, but as we show this can be ambiguous. More specifically, we obtain the following results: We introduce a new dissimilarity measure *d*_*LGT*_ and show that it defines a metric space on the set of *all* LGT networks with the same leaf taxa. Our measure can also be extended to tree-based networks.

The metric is based on operations to transform one network into the other, and so it can be used to explore the space of all LGT networks on the same taxa (which is useful for stochastic and hill-climbing reconstruction methods that make local moves on networks, the currently dominant approach [4,5]). We obtain a complexity dichotomy: *d*_*LGT*_ can be computed in linear time if the order of transfers along a species branch does not matter, and NP-hard if it does. In the latter case, we show that *d*_*LGT*_ is fixed-parameter tractable (FPT) in the *level* (maximum number of reticulations in a biconnected component) of the input networks. We implemented our algorithms and perform three sets of experiments. First, we show on random simulated networks that our implementation is usable in practice for large networks (up to ∼ 1800 vertices). We then show its usefulness in comparing LGT network predictions in two proofs-of-concept. In the first one, we compute *d*_*LGT*_ values between the outputs of the character-based methods presented in [34] and show quantitatively that transfer highway predictions are highly dependent on the approach and parameters used. In the other application, we look at networks predicted with a reconciliation tool (Ranger-DTL [31]), and show how our metric can be used to tune cost values within the reconciliation. Due to space constraints, some of the content (proofs, pseudo-code…) has been deferred to the Appendix, or is only present in the full online version (https://www.biorxiv.org/content/10.1101/2025.11.20.689557).

## 2 LGT networks and Tree-based networks

We first introduce all mathematical notions required to define our metric. The terminology is based on the definition of LGT networks presented in [28,38]. In a directed graph, the in-neighbors of a vertex are called its *parents* and its out-neighbors are *children*. A *phylogenetic network*, or just *network* for short, is a *directed acyclic graph* that has a single vertex of indegree 0, called its *root*. We also require that each vertex of outdegree 0, called a *leaf*, has indegree 1. Note that we allow vertices of indegree 1 and outdegree 1, which we call *subdivision vertices*. Given a phylogenetic network *N*, we denote its set of vertices by *V* (*N*), its set of arcs by *E*(*N*), and its set of leaves by *L*(*N*). The *contraction* of an arc (*u, v*) of a network *N* is an operation that identifies *u* and *v*, that is, it: adds the arc (*w, u*) for each in-neighbor *w* of *v*; adds the arc (*u, w*) for each out-neighbor *w* of *v*; removes *v* and its incident arcs. If *v* is a vertex with a single parent, contracting *v* means contracting the only arc entering *v*.

Two networks *N*_1_, *N*_2_ are *isomorphic*, denoted *N*_1_ ≃ *N*_2_, if *L*(*N*_1_) = *L*(*N*_2_) and there exists a bijective function *ϕ* : *V* (*N*_1_) → *V* (*N*_2_) such that: (1) for each *ℓ* ∈ *L*(*N*_1_), *ϕ*(*ℓ*) = *ℓ*; (2) (*u, v*) ∈ *E*(*N*_1_) if and only if (*ϕ*(*u*), *ϕ*(*v*)) ∈ *E*(*N*_2_). Furthermore, the two networks are *homeomorphic* if they are isomorphic when we ignore subdivision vertices. That is, *N*_1_ and *N*_2_ are homeomorphic if 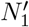 and 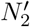 are isomorphic, where 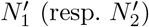 is obtained from *N*_1_ (resp. *N*_2_) by contracting each subdivision vertex.

### Trees and clusters

A *tree T* is a special type of network in which the underlying undirected graph has no cycle. For a vertex *v* of *T*, we write *L*_*T*_ (*v*) for the set of leaves descending from *v*, which we call the *cluster* of *v* in *T*. Note that two trees are isomorphic if and only if they have the same clusters, and each cluster occurs with the same multiplicity in both trees (recall that we allow subdivision vertices). Two trees are homeomorphic if and only if their sets of clusters are the same, without considering multiplicities.

### LGT networks

An *LGT network* is a pair 𝒩 = (*N*, (*E*_*p*_, *E*_*t*_)) where *N* = (*V, E*) is a network, (*E*_*p*_, *E*_*t*_) is a pair of subsets of *E* such that *E*_*p*_ ∪ *E*_*t*_ = *E* and *E*_*p*_ ∩ *E*_*t*_ = ∅, with *E*_*p*_ the *principal arc set* and *E*_*t*_ the *transfer arc set*; and the subgraph *T*_0_(𝒩) = (*V, E*_*p*_) consisting only of principal arcs is a tree with the same set of leaves as *N*. The tree *T*_0_(𝒩) is called the *base tree* of 𝒩 (note that *T*_0_(𝒩) may contain subdivision vertices, see Figure 1). A vertex with at least two children in *T*_0_(𝒩) is called a *tree vertex* (of either *N* or *T*_0_(𝒩)). Moreover, a vertex of *N* (or *T*_0_(𝒩)) that is incident to a transfer arc in *N* is called an *attachment point* (named as such since that endpoint that exists to attach a transfer). Since one goal of LGT networks is to distinguish between vertical and horizontal evolution, we assume the following.

**Fig. 1:**
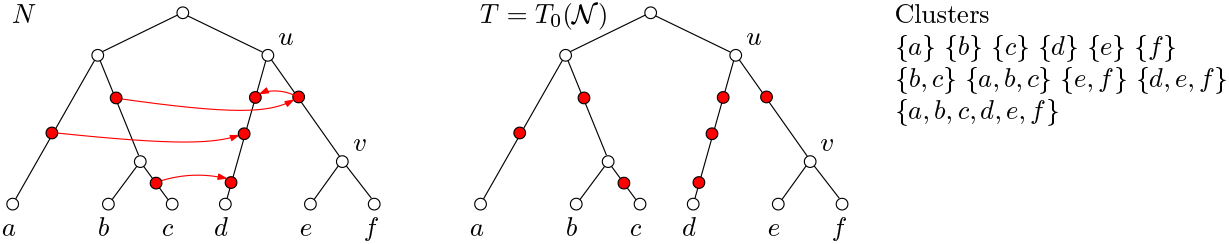
Left is an LGT network 𝒩= *N*|*T*. The base tree *T* = *T*_0_(𝒩) is shown in the middle. In *N*, black arcs are those of the base tree *T* (their direction is omitted, they point downwards) and red arcs are transfer arcs. White vertices are tree vertices and filled red vertices are attachment points. The set of clusters of *T* is shown on the right. Multiplicities are ignored, but for instance {*d*} occurs four times and {*e, f*} two times.

#### Assumption 1

*In an LGT network* 𝒩, *all attachment points are subdivision vertices in T*_0_(𝒩).

The example in Figure 1 satisfies this assumption. Note, this is not required in the original definition of LGT networks. It always holds in binary LGT networks, but our networks are not necessarily binary.

It will often be more useful to refer to the base tree directly instead of using the set of arcs *E*_*p*_ and the tree *T*_0_. We therefore use the notation *N*|*T* to denote the unique LGT network 𝒩whose network is *N* and whose base tree *T*_0_(𝒩) is *T*, and stick with this alternate notation for the rest of the paper. We sometimes use the calligraphic 𝒩 to denote an LGT network without specifying *N* and *T*. Note that the *N*|*T* notation lets us deduce the arc partition of 𝒩 with *E*_*p*_ = *E*(*T*) and *E*_*t*_ = *E*(*N*) \ *E*(*T*). Using this notation, given an LGT network *N*|*T*, we write *v*⪯_*T*_ *u* to indicate that *v* is a descendant of *u* in *T*.

A *tree pair* in *N*|*T* is an ordered pair of vertices from *V* (*N*), denoted [*u, v*], such that *u, v* are tree vertices, *v* ≺_*T*_ *u*, and the directed path from *u* to *v* in *T* contains only attachment points, except *u* and *v* (we use brackets to distinguish tree pairs from arcs of *T*). Intuitively, a tree pair is just a branch linking tree vertices between which transfers occurred. Any attachment point on the *u* − *v* path is said to be *on* [*u, v*]. In Figure 1, [*u, v*] forms a tree pair, as well as [*u, d*]. Finally, we need a notion of identical LGT networks, since isomorphims may not preserve principal and transfer arcs. We say that two LGT networks 𝒩_1_ = *N*_1_|*T*_1_, 𝒩_2_ = *N*_2_|*T*_2_ are *LGT-isomorphic*, denoted 𝒩_1_ ≃ _*LGT*_ 𝒩_2_, if there exists an isomorphism *ϕ* between *N*_1_ and *N*_2_ such that *ϕ* is also an isomorphism between *T*_1_ and *T*_2_.

### Tree-based networks

A network *N* = (*V, E*) is a *tree-based network* if there *exists* a pair (*E*_*p*_, *E*_*s*_) such that (*N*, (*E*_*p*_, *E*_*s*_)) is an LGT network with the same leaves as *N*. In this case, (*V, E*_*p*_) is called a *base tree* of *N*. This is equivalent to asking for a subset of arcs *E*_*p*_ such that (*V, E*_*p*_) is a tree with the same set of leaves as *N* [35]. The main difference is that in an LGT network, the base tree is given, whereas a tree-based network may contain many base trees. The set of all base trees of a tree-based network *N* will be denoted *B*(*N*). Thus, the set of all LGT networks contained in a tree-based network *N* is {*N*|*T* : *T* ∈ *B*(*N*)}.

### Time consistent LGT networks

We say that an LGT network *N*|*T* is *timeconsistent* if their exists a labeling function *λ* : *V* (*N*) → ℕ^+^ such that: (1) for any transfer arc (*u, v*), *λ*(*u*) = *λ*(*v*); and (2) for any other arc (*u, v*), *λ*(*u*) *< λ*(*v*). In other words, *λ* represents time that increases from past to present. The arcs of the base tree go in strictly increasing times, as they represent vertical evolution, whereas arcs representing horizontal evolution require co-existence and thus have the same time. The LGT network in Figure 1 is easily seen to be time-consistent.

## 3 A metric for LGT networks and tree-based extensions

Given two LGT networks on the same leaves, we want to compare both their base trees and their transfer arcs. For the latter, we delete transfers to eliminate disagreements. As for base trees, they can be compared using established tree metrics, and we here use Robinson-Foulds. To avoid situations where nonattachment points become incident to transfer arcs, we only allow contraction of arcs whose two ends are not attachment points. Given an LGT network *N*|*T*, we thus define the two following operations (see Figure 2):

**Fig. 2:**
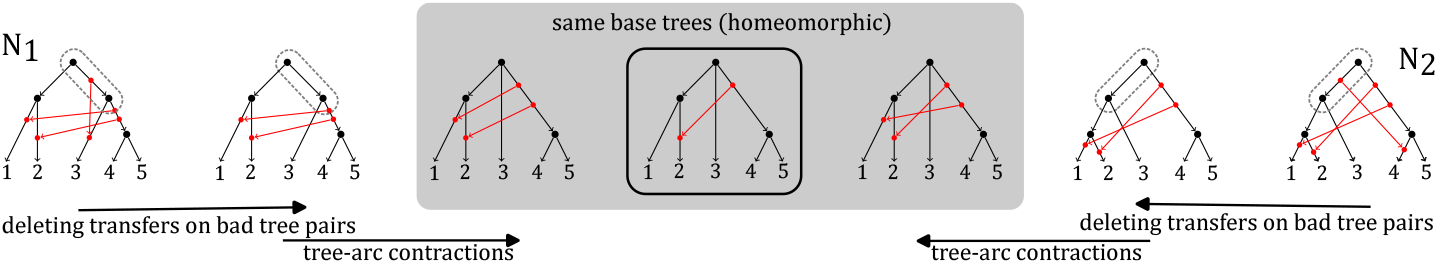
Illustration of the different steps of an edition pathway between two networks 𝒩_1_, 𝒩_2_ (leftmost and rightmost networks). We want to contract “bad” tree pairs (encircled in 𝒩_1_ and 𝒩_2_, which respectively correspond to bad clusters 3, 4, 5 and 1, 2, 3). Because tree-arc contractions forbid attachment points, we must first delete the transfer arcs on those tree-pairs. Once bad tree pairs are removed, the base trees are homeomorphic (grey area), and one must then compute *d*_*TR*_. As a side note, here 𝒩_1_ is not time-consistent because of the vertical transfer arc.

– the *contraction* of an arc (*u, v*) of *E*(*N*), where *u* and *v* are both non-leaf tree vertices of *N*|*T*. Since *u* and *v* are not incident to a transfer, (*u, v*) is also in *E*(*T*). This operation results in another LGT network *N*′|*T*′, where *N*′ (resp. *T*′) is obtained by contracting (*u, v*) in *N* (resp. in *T*);
– the *deletion* of a transfer arc (*u, v*) of *E*(*N*) \ *E*(*T*). This operation results in another LGT network *N*′|*T*′ where *N*′ is obtained by removing arc (*u, v*) from *N*, and contracting any resulting subdivision vertices of *N* (note, only *u* or *v* can become subdivision vertices); *T*′ is obtained from *T* by contracting the same subdivision vertices that were contracted in *N*.

An LGT network *N*′|*T*′ is an *LGT reduction* of *N*|*T* if *N*′|*T*′ can be obtained from *N*|*T* by applying a sequence of contractions and transfer deletions, as defined above. We define *δ*(*N*|*T, N*′|*T*′) as the number of operations required to transform *N*|*T* into *N*′|*T*′. Given two LGT networks 𝒩_1_ = *N*_1_|*T*_1_, 𝒩_2_ = *N*_2_|*T*_2_, we say that *N*|*T* is a *common LGT reduction* of 𝒩_1_, 𝒩_2_ if *N*|*T* is LGT-isomorphic to an LGT reduction of both 𝒩_1_ and 𝒩_2_. The LGT distance is:

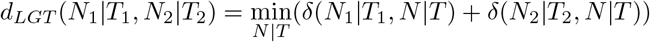

where the minimization is over all common LGT reductions *N*|*T*. When such an *N*|*T* minimizes the above expression, it is called a *maximum common LGT reduction*. See Figure 2.

### Remark on exploring spaces of LGT networks

A common LGT reduction always exists if the leaf sets are the same, since on both networks we can delete every transfer arc, resulting in a tree, and then contract all remaining arcs to obtain a star tree. By defining the reverse of contractions and transfer deletions, one could therefore transform any LGT network into another (as long as leaves are the same). Thus our operations and their reverse can be used to traverse the space of all LGT networks, as claimed in the introduction.

### Extending to tree-based networks

Note, *d*_*LGT*_ translates naturally to tree-based networks, by asking for the base trees that minimize *d*_*LGT*_. That is, if *N*_1_ and *N*_2_ are tree-based networks, the distance is:

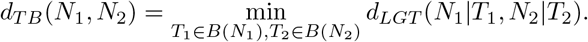

### 3.1 A special case: the transfer reduction distance

A special case of *d*_*LGT*_ occurs when the base trees of the given LGT networks are the same (i.e., homeomorphic), and all disagreements are just between transfer arcs. An LGT network *N*′|*T*′ is a *transfer reduction* of *N*|*T* if it is obtained from *N*|*T* by applying a sequence of transfer deletions only. Observe that the number of transfer deletions needed to transform *N*|*T* into *N*′|*T*′ is|*E*(*N*) \ *E*(*T*)| −|*E*(*N*′) \ *E*(*T*′)|. We say that *N*|*T* is a *common transfer reduction* of two LGT networks *N*_1_|*T*_1_ and *N*_2_|*T*_2_ if *N*|*T* is LGT-isomorphic to a transfer reduction of both *N*_1_|*T*_1_ and *N*_2_|*T*_2_ We can define the *transfer reduction* distance as

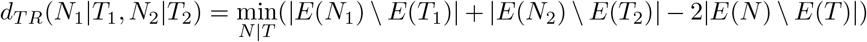

where the minimization is over all common transfer reductions *N*|*T*. An *N*|*T* that minimizes this expression is called a *maximum common transfer reduction*. Note that a common transfer reduction does not always exist, and in fact it exists if and only if *T*_1_ and *T*_2_ are homeomorphic.

### 3.2 Useful properties of *d*_*LGT*_

We show that in order to compute *d*_*LGT*_, it suffices to compare the base tree and the transfer arcs separately. We first define a weighted RF (*wRF*) distance for the tree part. Let 𝒩_1_ = *N*_1_|*T*_1_ and 𝒩_2_ = *N*_2_|*T*_2_ be two LGT networks. A vertex *v* of *N*_1_ is *bad* (w.r.t. 𝒩_1_ and 𝒩_2_) if *v* is a tree vertex and the cluster *L*_*T*_ (*v*) is not a cluster of *T*_2_. Likewise, a vertex *v* of *N*_2_ is bad if it is a tree vertex and 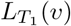 is not a cluster of *T*_1_. Now let *v* be a non-root tree vertex of either network, and let *u* be the unique tree vertex such that [*u, v*] is a tree-pair (recall, tree-pairs have only attachment points between them). Assign the weight *w*(*v*) as the number of transfer arcs that have an endpoint on [*u, v*], plus one. Then the *wRF* distance is defined as the sum of weights of bad vertices, i.e.,

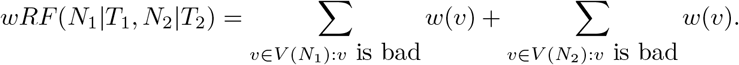

In Figure 2, each network has a bad vertex of weight two, so *wRF* (𝒩_1_, 𝒩_2_) = 4. For the transfer component, let (*u, v*) be a transfer arc of *N*_1_|*T*_1_. Let [*u*_1_, *u*_2_] be the unique tree pair that *u* is on, and let [*v*_1_, *v*_2_] be the unique tree pair that *v* is on. We say that (*u, v*) is a *bad transfer* if one of *u*_2_ or *v*_2_ is a bad vertex. We also say that (*u, v*) is a *doubly bad transfer* if both *u*_2_, *v*_2_ are bad. Apply the analogous definitions to the transfers of *N*_2_|*T*_2_. In Figure 2, each network has one bad transfer, but no doubly bad ones.

#### Theorem 1

*Given two LGT networks* 𝒩_1_ = *N*_1_|*T*_1_, 𝒩_2_ = *N*_2_|*T*_2_ *on the same leafsets*,

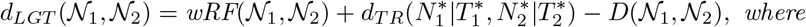

– 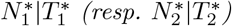 *is the LGT network obtained from N*_1_|*T*_1_ *(resp. N*_2_|*T*_2_*) by applying a transfer deletion on each of its bad transfers (including doubly bad ones), and then contracting every bad vertex*.
– *D*(𝒩_1_, 𝒩_2_) *is the number of doubly bad transfer arcs that are in* 𝒩_1_ *or in* 𝒩_2_.

In essence, *wRF* counts the number of bad transfers and tree-arc contractions that must be performed, and *d*_*TR*_ adds the transfer deletions needed after (*wRF* double-counts doubly bad transfers so *D*(𝒩_1_, 𝒩_2_) is subtracted). Day’s algorithm can be used to compute *wRF* in linear time [39]. The core of the problem is therefore to compute *d*_*TR*_. We conclude this section by showing that *d*_*LGT*_ is a true mathematical metric. Recall that this requires showing that: (1) *d*_*LGT*_ (𝒩_1_, 𝒩_2_) = 0 if and only if the two networks are LGT-isomorphic; (2) the distance is symmetric; and (3) it satisfies the triangle inequality.

#### Theorem 2

*The d*_*LGT*_ *distance satisfies all conditions of a metric*.

## 4 The complexity of computing *d*_*T R*_ (and *d*_*LGT*_)

We show that there is a dichotomy in the complexity of computing *d*_*TR*_, depending on whether we care about the order of transfers along a tree pair. That is, suppose we have an LGT network *N*|*T* and that there is a tree pair [*u, v*] with more than one attachment points on it. This assumes an ordering of the transfers that occurred between *u* and *v*. However, in many cases, this order is unknown and very difficult to predict accurately, and it may be sufficient to contract all these attachment points into one due to uncertainty. This leads to LGT networks with at most one attachment point per tree pair. This matters, because we can compute *d*_*TR*_ in linear time under this assumption, and otherwise computing *d*_*TR*_ is NP-hard. This is true even if there are at most 3 attachment points on tree pairs, leaving the case of 2 attachment points open. Unless stated otherwise, we now deal with pairs of LGT networks whose base trees are homeomorphic. To simplify notation, we shall assume that two networks have the same set of tree vertices (instead of relying on an homeomorphism). Thus for two LGT networks *N*_1_|*T*_1_, *N*_2_|*T*_2_, we may assume that [*u, v*] is a tree pair of *T*_1_ if and only if [*u, v*] is a tree pair of *T*_2_ (though the number of attachment points may vary).

### 4.1 One attachment point per tree pair

Let *N*|*T* be an LGT network, possibly non-binary. We define the set of tree pairs that share the two ends of a transfer, as follows:

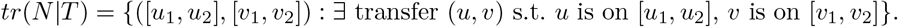

In words, ([*u*_1_, *u*_2_], [*v*_1_, *v*_2_]) ∈ *tr*(*N*|*T*) means that some transfer has its tail between *u*_1_ and *u*_2_, and its head between *v*_1_ and *v*_2_. Note, *tr*(*N*|*T*) contains ordered pairs (of tree pairs), and ([*v*_1_, *v*_2_], [*u*_1_, *u*_2_]) is different. Since we assume that tree vertices of 𝒩_1_ and 𝒩_2_ are the same, we can compare *tr*(𝒩_1_) and *tr*(𝒩_2_) directly. When there is at most one attachment point per tree pair, it suffices to delete transfers whose corresponding pair in the *tr* set is not in the other. We can thus use Day’s algorithm for the *wRF* part [39], and compute the symmetric difference for the transfer part. Recall that *A*Δ*B* = (*A* \ *B*) ∪ (*B* \ *A*).

#### Theorem 3

*Let* 𝒩_1_ = *N*_1_|*T*_1_, 𝒩_2_ = *N*_2_|*T*_2_ *be two LGT networks such that T*_1_, *T*_2_ *are homeomorphic. Suppose that for any tree pair* [*u, v*] *of either network, at most one attachment point is on* [*u, v*]. *Then:*

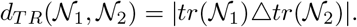

*Consequently, d*_*LGT*_ (𝒩_1_, 𝒩_2_) *can be computed in time O*(*m*_1_ + *m*_2_), *where m*_1_ = |*E*(*N*_1_)|, *m*_2_ = |*E*(*N*_2_)|.

Beyond one attachment point, the symmetric difference *tr*(𝒩_1_) Δ *tr*(𝒩_2_) may fail because the order may disagree. Figure 3 illustrates this phenomenon, which is exploited in the NP-hardness proof that follows.

**Fig. 3:**
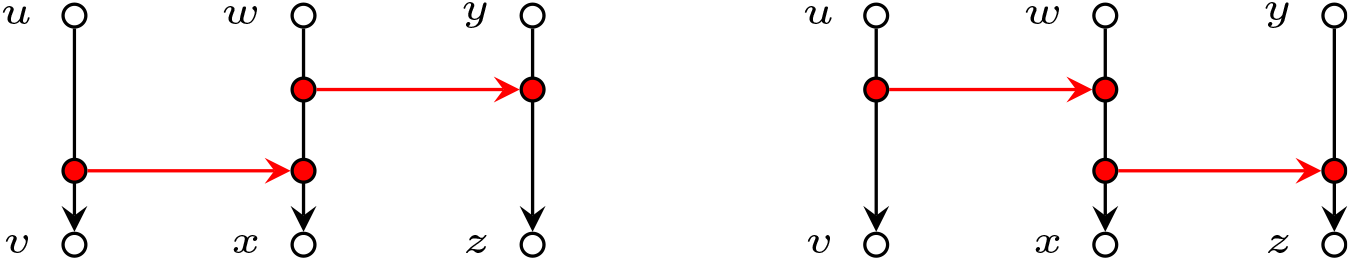
With more than one attachment point per tree pair, one may have *tr*(𝒩_1_) = *tr*(𝒩_2_) even though the networks are different. We have *tr*(𝒩_1_) = *tr*(𝒩_2_) = {([*u, v*], [*w, x*]), ([*w, x*], [*y, z*])}, and yet the two LGT networks cannot be isomorphic.

### 4.2 Three attachment points, when the order of transfers matters

We show that computing *d*_*LGT*_ is NP-hard, even if the input is restricted to time-consistent networks. The hardness arises when many transfers are in reverse order as in Figure 3. Such pairs of transfers are incompatible and we can keep at most one in each network. When many such incompatibilities arise, we have to find a maximum number of pairwise-compatible transfers, a difficult problem. Inspired by [40], our reduction is from 3-SAT, so that maximally compatible transfers correspond to satisfying a formula.

#### The reduction (see Figure 4)

In 3-SAT, we receive a boolean formula *ϕ* over variables *x*_1_, …, *x*_*n*_, with *ϕ* a conjunction of clauses *c*_1_, …, *c*_*m*_, each containing three literals (a literal is a variable *x* or its negation 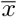). We must decide if there is a variable assignment that makes every clause *true*. From *ϕ*, we construct time-consistent LGT networks *N*_1_|*T*_1_, *N*_2_|*T*_2_ and an integer *k*, such that *ϕ* is satisfiable ⇔ *d*_*TR*_ (*N*_1_|*T*_2_, *N*_1_|*T*_2_) ≤ *k*.

**Fig. 4:**
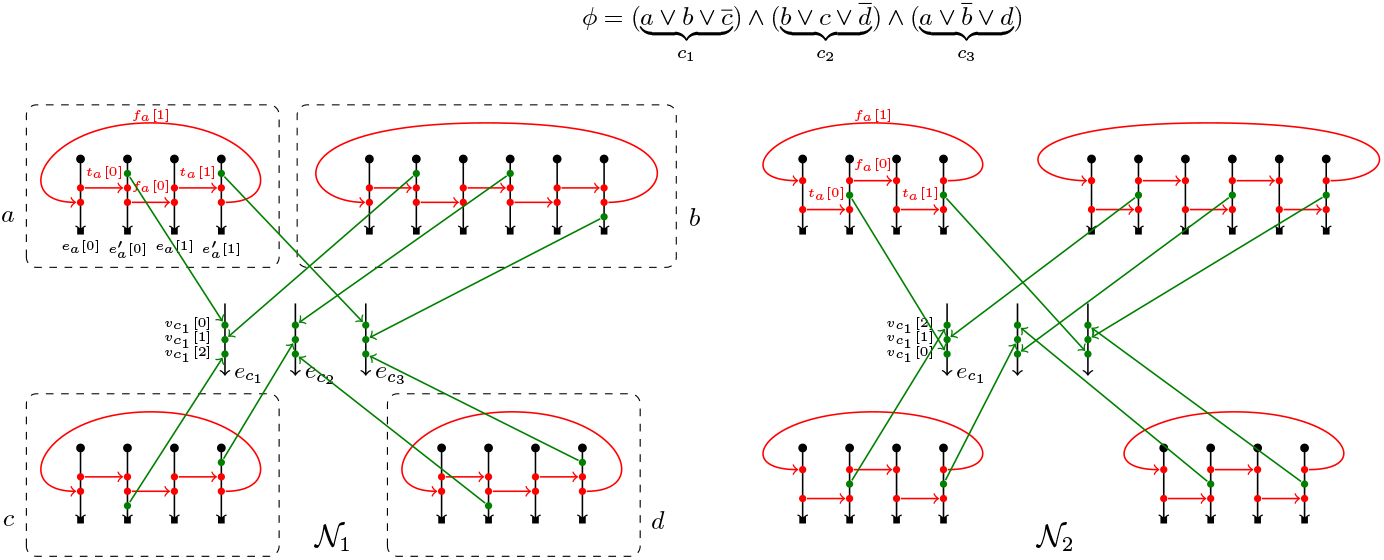
Illustration of the reduction used in the proof of Theorem 4 (NP-completeness of *d*_*LGT*_).

For a variable *x*, we write *d*(*x*) for the number of clauses in which *x* appears. We denote the clauses that contain *x* as *c*_*x*,0_, *c*_*x*,1_, …, *c*_*x,d*(*x*)−1_, ordered arbitrarily. To construct *N*_1_|*T*_1_ and *N*_2_|*T*_2_, the any base tree will do, so we let *T* be any tree with 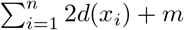 leaves. Denote by *E* the set of arcs of *T* whose head is a leaf. To obtain *T*_1_ and *T*_2_, we start from *T* and add attachment points only to arcs in *E* (this is for time-consistency). The arcs in *E* will therefore become tree pairs. Given a transfer arc *t* = (*u, v*), we denote *t*.tail = *u* and *t*.head = *v*.

#### Variable gadget

We describe variable gadgets in *N*_1_|*T*_1_. The construction of the base tree starts from *T*. For each variable *x*, choose 2*d*(*x*) arcs in *E* that we denote 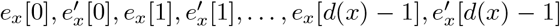 (note, each variable has its own distinct arcs). Subdivide each of these arcs twice to create two attachment points per arc. For *i* ∈ [0; *d*(*x*) − 1], we introduce a transfer arc *t*_*x*_[*i*] from the highest attachment point on *e*_*x*_[*i*] to the highest attachment point on 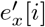. These represent assigning *x* to true. Then, for all *i* ∈ [0; *d*(*x*) − 1], add a transfer arc *f*_*x*_[*i*] from the lowest attachment point on 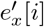 to the lowest attachment point on *e*_*x*_[*i* + 1] (we take *i* + 1 modulo *d*(*x*)). In *N*_2_|*T*_2_, we do the same, except that we **reverse** the order of attachment points of *t*_*x*_[*i*] and *f*_*x*_[*i*]. That is, the *f*_*x*_[*i*]’s take the highest attachment points from 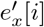 to *e*_*x*_[*i* + 1], and the *t*_*x*_[*i*]’s take the lowest from *e*_*x*_[*i*] to 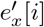.

#### Clause gadget

For each clause *c* of *ϕ*, we denote the variables that occur in *c* as *x*_*c*,0_, *x*_*c*,1_, *x*_*c*,2_, with the only constraint that in this ordering, all positive literals appear before all negative literals. Let us now get back to modifying *N*_1_|*T*_1_. For each clause *c*, choose an arc *e*_*c*_ of *E* that is not used for a variable gadget or for another clause gadget. Add three attachment points *v*_*c*_[0], *v*_*c*_[1], *v*_*c*_[2] on *e*_*c*_, in this order from top to bottom. For *j* ∈ {0, 1, 2}, consider the *j*-th variable *x* := *x*_*c,j*_. Let *i* be the rank of *c* among the clauses in which *x* appears, i.e., *c* = *c*_*x,i*_. If *x* appears as a positive literal in *c*, we add an attachment point *a*_*c,x*_ above *t*_*x*_[*i*].head on 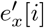. If *x* appears as a negative literal in *c, a*_*c,x*_ is added below *f*_*x*_[*i*].tail on 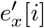. We add a transfer arc from *a*_*c,x*_ to *v*_*c*_[*j*]. In *N*_2_|*T*_2_, add attachment points *v*_*c*_[2], *v*_*c*_[1], *v*_*c*_[0] on *e*_*c*_ (so, in reverse order). For *j* ∈ {0, 1, 2} and variable *x* := *x*_*c,j*_, let *i* be such that *c* = *c*_*x,i*_. Add attachment point *a*_*c,x*_ on 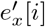 above *t*_*x*_[*i*].head and below *f*_*x*_[*i*].tail (whether *x* is positive or not). Then add a transfer arc from *a*_*c,x*_ to *v*_*c*_[*j*].

We denote 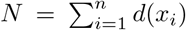. We show that *ϕ* is satisfiable if and only if *d*_*TR*_(*N*_1_|*T*_1_, *N*_2_|*T*_2_) ≤ 2*N* + 4*m*.

##### Theorem 4

*Computing d*_*LGT*_ *is NP-complete, even on binary time-consistent networks with at most 3 transfers per attachment point*.

*Proof (sketch)*. Time-consistency can be established by providing an explicit time-labeling, see Appendix. For the equivalence with the satisfiability problem, first assume that *ϕ* is satisfied by some assignment. If variable *x* is assigned true, then we delete all *f*_*x*_[*i*] arcs and keep only the *t*_*x*_[*i*] arcs, in both networks. Do the opposite if *x* is false. This amounts to 2*N* deletions (*N* per network). Then for every clause *c*, take a variable *x* that satisfies it, and on both networks delete every transfer arc that enters *e*_*c*_, except the arc from *a*_*c,x*_ to *v*_*c*_[*j*] (where *j* ∈ {0, 1, 2}). Thus for each clause we perform two transfer deletions, which amounts to 2*m* deletions per network. The total is 2*N* +4*m*, as desired. One can see in the Appendix (Figure 9) that doing these deletions on both networks results in the same transfer reduction. In the other direction, assume that *d*(*N*_1_|*T*_1_, *N*_2_|*T*_2_) ≤ 2*N* + 4*m*. One can argue that any solution must essentially mimic the previous direction. In an *x* gadget, we cannot keep both a *t*_*x*_[*i*] and *f*_*x*_[*i*] arc because their orders are reversed in the two networks. The best we can do is keep all *t*_*x*_[*i*] or all *f*_*x*_[*i*] arcs. The ones we keep correspond to an assignment of *x*. In a *c* gadget, we can keep at most one transfer arc going on *e*_*c*_ because the order of attachment points are reversed in the two networks. To perform at most 2*N* + 4*m* deletions, one such arc must be kept, which corresponds to the variable that can satisfy *c*.

This reduction can also be adapted to prove the NP-hardness of *d*_*TB*_, the distance on tree-based networks. The idea is to add a gadget to *N*_1_ and *N*_2_ to constraint their base trees, such that transfer arcs are the same.

##### Theorem 5

*Computing d*_*TB*_ *is NP-hard*.

### 4.3 Parameterized algorithms and pre-processing rules

We first observe that to compute *d*_*TR*_ in practice, and thus *d*_*LGT*_, one can design a simple pre-processing rule as follows.

#### Transfer cleaning rule

Given two networks 𝒩_1_ = *N*_1_|*T*_1_ and 𝒩_2_ = *N*_2_|*T*_2_, with *T*_1_, *T*_2_ homeomorphic, if 𝒩_1_ has a transfer from tree pair [*a, b*] to tree pair [*c, d*], but 𝒩_2_ has no such transfer, then we delete this transfer immediately (and remember to count it in the distance).

The transfer cleaning rule is easily seen to be correct by observing that no edge contraction nor transfer deletion may introduce new transfer arcs between tree pairs of 𝒩_2_. We note that this pre-processing rule is critical in achieving the practical scalability results presented in the experiments section. After cleaning, we can test all ways of deleting transfers.

##### Proposition 1

*The distance d*_*LGT*_ (𝒩_1_, 𝒩_2_) *can be computed in time O*(2^*t*^ · *m*), *where t (resp. m) is the number of transfer arcs (resp. arcs) in* 𝒩_1_ *and* 𝒩_2_.

On binary networks, we can do better by handling matching blobs in the two networks independently (see appendix and [11] for a definition of the level).

##### Theorem 6

*Given two binary LGT networks* 𝒩_1_, 𝒩_2_ *both of level at most ℓ, one can compute d*_*TR*_(𝒩_1_, 𝒩_2_) *and thus d*_*LGT*_ (𝒩_1_, 𝒩_2_) *in time O*(4^*ℓ*^ · *m*^2^), *where m is the total number of edges in both networks*.

## 5 Experiments

All of the experiments use an implementation of the algorithms presented in Section 4.1 and 4.2, publicly-available at https://github.com/bmarchand/transfer_edition_distance. The algorithms were implemented in Rust and are available in Python with PyO3 (https://github.com/PyO3/pyo3).

We remark that for the first experiment, other measures designed to compare phylogenetic networks (reviewed in the introduction [9,10,11,12,13,14,15,16,17]) are not applicable, as the task is specifically to compare LGT networks, in which a distinction between tree arcs and transfer arcs is made. To illustrate the problem, consider the fact that a given tree-based network *N* may give rise to two different LGT networks *N*|*T* and *N*|*T*′ if *T, T*′ are two different base trees. The metrics mentioned above do not distinguish these situations. While this argument does not apply to the second and third experiment, we note that in both of them, the networks predicted by the methods are tree-based, but not necessarily orchard networks [26] (see Figure 12 in Appendix for more details). To our knowledge, none of the existing implemented metrics [17,3] are capable of distinguishing all tree-based networks.

### 5.1 Benchmarks on random LGT networks

#### Overview

We first evaluate the computational scalability of our metric on random LGT networks, using the simulator from [38]. It outputs binary LGT networks, potentially with several transfers along a tree-pair, with their orders specified. Our results indicate that, although *d*_*LGT*_ is NP-hard to compute, Algorithm 1 (Appendix) is fast in practice up to very large network sizes. Indeed, Figure 5 shows that under reasonable parameter choices, two output networks of size ≃ 1800 may be compared in ≲ 0.1 seconds.

**Fig. 5:**
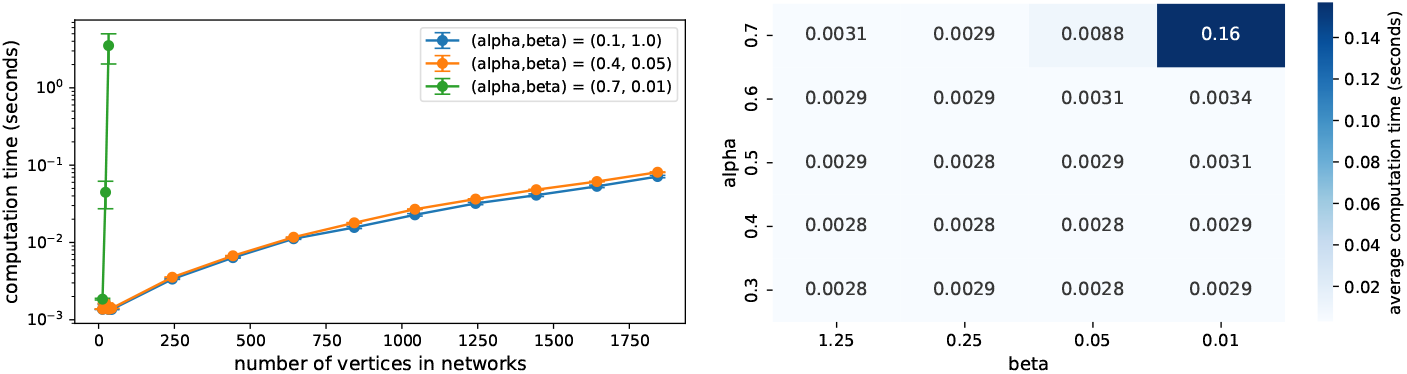
Run-time benchmark of the computation of our metric on random LGT networks.(left) For several combinations of (*α, β*), average computation time of the distance between two random networks, as a function of the number of vertices. (right) For a fixed network size (200), table giving the average computation time of the distance between two random networks, for various combinations of (*α, β*). Note that *α* = 0.7 is higher than values explored in [38] and arguably leads to unrealistic networks.

#### Simulator

The simulator in [38] takes three parameters: the “number of steps” *N* (each step adds two vertices to the network, see below), a transfer probability *α*, and a “level coefficient” *β*. Starting with a rooted tree with two leaves, at each step a random choice is made between a *speciation* and a *transfer*, with *α* the probability of a transfer. If “speciation” is picked, a leaf is chosen uniformly at random and a new leaf is attached on its parent arc. If “transfer” is picked, then two leaves *l* and *l*′ are chosen and a transfer is added on their parent arcs. Parameter *β* influences the choice of *l* and *l*′, with a higher *β* increasing the chances of choosing two leaves in different biconnected components. In essence, a higher *α* increases the number of transfers, and *β* controls the *level* (the maximum number of reticulations in a biconnected component).

#### Results

Figure 5 shows that our algorithm is fast on a wide range of parameters, except when *α* = 0.7 and *β* = 0.01. This can be seen both on Figure 5 (a), where (*α, β*) = (0.7, 0.01) exhibits a stronger exponential scalin, and on Figure 5 (b) for a fixed network size. We note that *α* = 0.7 is beyond the range of values explored in [38], and may be considered unrealistically high (recall that *α* = 0.7 means that, in expectation, 70% of events are transfers, and 30% speciations). Even then, hard instances are only produced if *β* ≤ 0.01, i.e., when transfers are 100× more likely between two leaves whose parents are in the same biconnected component, producing “blobs” with exceedingly many transfers.

### 5.2 Comparison of character-based transfer reconstructions

Some models such as *perfect phylogenetic networks* [41] and *perfect transfer networks* [33,42] can predict transfers on a species tree by explaining the presence/absence of given characters at the leaves (here, characters could be genes, functions, traits, …). We revisit the experiments and data used in one of the latest iterations of this line of work [34] to illustrate a use-case of *d*_*LGT*_. The authors of [34] were only able to compare their predictions qualitatively, and here our metric may be used to compare predictions *quantitatively*.

#### Overview

Given a species tree and characters at the leaves, [34] considers three different approaches for reconstructing the history of individual characters. For each character, they assume a *single origin* and *losses* and *transfers* as evolutionary events. A first approach is called *Basic labeling*, and infers a character at an internal vertex if and only if it is present in all of its descendants. The assumption is then that maximal subtrees containing a character are involved in an HGT. Such a labeling has sometimes been used in hypothesis-testing, where biologists want to check whether a gene could have emerged in a de novo fashion, or was rather acquired through HGT (see [43,44]). The two other approaches, *Genesis* and *Sankoff*, are more sophisticated dynamic programming schemes computing ancestral assignments of the characters minimizing the costs of losses and transfers. Details can be found in [34]. For each prediction method and each character, the set of inferred transfers yields an LGT network with the species tree as base tree (plus subdivisions). Note that the inferred transfers along a tree pair are *unordered*, so we may use Theorem 3.

#### Data

We use the same data as in [34]. It consists of a set of 180 functional characters from the KEGG database (Kyoto Encyclopedia of Genes and Genomes [45]) within a set of 45 bacterial species. We take the species tree connecting these species from the NCBI Taxonomy Browser [46].

#### Results

In [34], reconstructed networks were combined into sets of “transfer highways”, that were then compared regardless of their weights (i.e. regardless of how many transfers use the highway). Having a metric such as *d*_*LGT*_ allows for a finer quantitative comparison. On Figure 6, we report computations of *d*_*LGT*_ values between network predictions reconstructed with each of the three methods. For each of the 180 KEGG characters mentioned above (x-axis on Figure 6), the reconstructions are compared in a pairwise fashion: Basic vs. Genesis, Basic vs. Sankoff, and Genesis vs. Sankoff (y-axis on Figure 6). We note that the methods “Genesis” and “Sankoff” yield predictions that are much closer to one another, compared to “Basic”. While this analysis does not tell which method is more accurate, it indicates that the choice of predictor significantly alters the results. Hence, care must be taken when making biological conclusions from HGT predictions as they may be highly dependent on the method used. Nevertheless, these results are an example of how a metric may be used to “cluster” predictions into similar groups, so that a consensus may be identified.

**Fig. 6:**
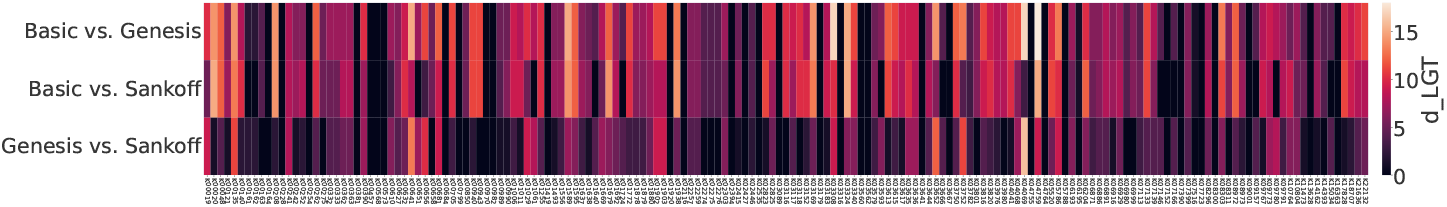
Matrix of distances between reconstructed phylogenetic networks, using the character-based methods presented in [34], dubbed “Basic”, “Sankoff” and “Genesis”. Each column of the matrix corresponds to a character. For each character, each method reconstructs a network. We observe that Genesis and Sankoff tend to give network reconstructions that are closer to one another, compared with Basic. In [34], only qualitative comparisons had been reported.

A full pairwise comparison of the networks predicted by each method for a subset of characters is also given in Appendix, Figure 13.

### 5.3 Optimizing reconciliation parameters for transfer prediction Overview

Reconciliation addresses the following problem: given a species tree *S* and a gene tree *G*, with a map from *L*(*G*) to *L*(*S*), find an evolutionary history on *G* that explains their differences, typically with duplication, loss, and transfer events. This is often done parsimoniously by minimizing the weights of inferred events. A crucial problem in this context is to know what cost to associate to each allowed type of event (see e.g. [48]). We show how metrics may help tune the parameters of reconciliation tools, to optimize for transfer inference.

#### Experimental Protocol

A set of *N* = 100 histories, each consisting of a species tree with *L* = 50 leaves and a gene tree evolving inside it were simulated with Asymmetree [47], using rates for duplications, losses, and transfers of respectively 0.5, 0.5 and 0.1 (see the documentation of Asymmetree for details). This simulator includes speciations, duplications, losses and transfers events. By adding arcs corresponding to the transfers in the simulation to the species tree, we get a “ground truth” network. We also give each pair of species tree and gene tree to RangerDTL [31], a leading tool for reconciliation. Its output is a mapping of each vertex of the gene tree into the species tree, along with a labeling with one of the events (speciation, duplication, transfer). We run RangerDTL *p* = 20 times per pair of species/gene trees, to get a variety of optimal reconciliations (there may be many). Likewise, by adding the transfers onto the species tree, we get a “reconstructed LGT network”, see Figure 7 (a). Note that both the ground truth and reconstructed networks are unordered, as described in Section 4.1, i.e. with at most one attachment point per tree pair. In the (rare) case that two transfer events take place between the same tree pair, we treat it as a single transfer. RangerDTL also supports cost values for losses, duplications and transfers (the default being 1,2 and 3). We fix the costs of duplications and losses in RangerDTL to 10, and use transfer costs ranging from 10 to 100. Figure 7 (b) shows normalized *d*_*LGT*_ values between the reconstructed and ground truth networks, as a function of the cost of transfers, averaged over *N* and *p*. species as base tree and (2) are unordered. Formally, for the normalization we compute 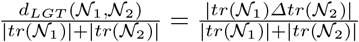.

**Fig. 7:**
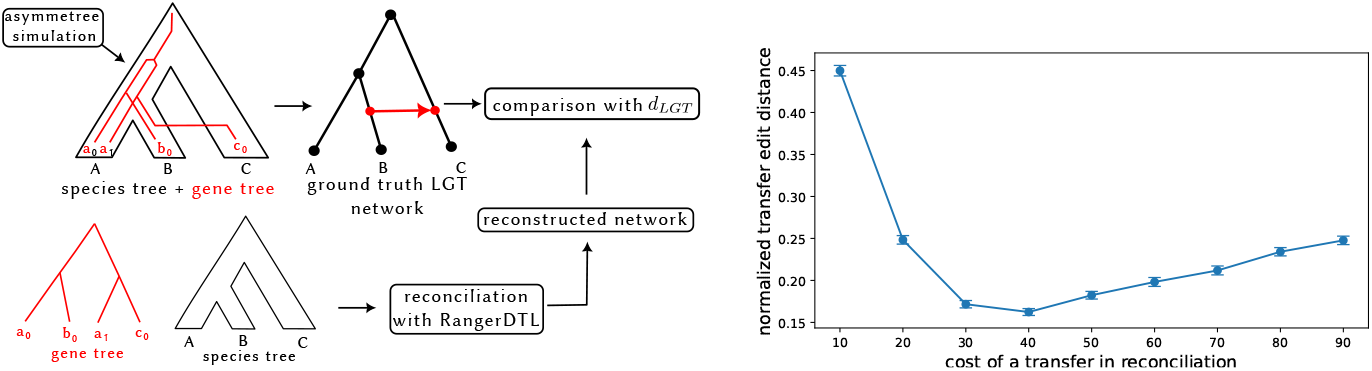
(left) A set of species tree along with a gene tree evolving within it (with duplications, transfers and losses), are simulated using Asymmetree [47]. Adding the transfers to the species tree yields a ground-truth LGT network. A reconciliation of each gene-tree/species-tree pair is also computed using RangerDTL [31], yielding a reconstructed LGT network. (right) Average normalized *d*_*LGT*_ values between the reconstructed and ground truth networks, as a function of the cost associated to transfer events in the reconciliation (the costs of duplications and losses is kept constant). This plot allows the identification of cost values that minimize the average reconstruction distance, offering a way to calibrate reconciliation costs.

## Results

Figure 7 (b) displays the normalized distance as a function of the transfer cost chosen in the reconciliation. We observe the presence of an optimal cost of 40, yielding reconstructions closest to the ground truth. Such a setup may be used to tune the parameters of RangerDTL (or other similar tools), prior to applications in cases where the ground truth is not known.

## 6 Discussion and Conclusion

Future research directions could include methods for computing the *consensus* of more than two networks, for instance through a *median* under our distance. One could also explore other popular choices of edit operations for the species tree, such as *subtree-prune and regraft* (SPR) or *nearest-neighbor interchange* (NNI). It should also be noted that the complexity of *d*_*TR*_ when tree pairs have at most 2 attachment points is open. Another algorithmic question of interest is whether computing our metric is fixed-parameter tractable in *d*_*LGT*_, e.g., with a complexity that is exponential in *d*_*LGT*_ only and not the sizes of the networks. To finish, one could also explore generalizations of our metric that would take into account the “distance” between disagreeing transfers, to better distinguish a slightly misplaced transfer from one further away.

## Appendix complete proofs

### A Proofs for Section 3 (A metric for LGT networks and tree-based extensions)

#### Theorem 1

*Given two LGT networks* 𝒩_1_ = *N*_1_|*T*_1_, 𝒩_2_ = *N*_2_|*T*_2_ *on the same leafsets*,

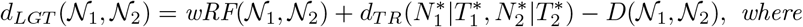

– 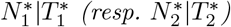 *is the LGT network obtained from N*_1_|*T*_1_ *(resp. N*_2_|*T*_2_*) by applying a transfer deletion on each of its bad transfers (including doubly bad ones), and then contracting every bad vertex*.
– *D*(𝒩_1_, 𝒩_2_) *is the number of doubly bad transfer arcs that are in* 𝒩_1_ *or in* 𝒩_2_.

*Proof*. First notice that for any LGT network *N*|*T*, by applying a transfer deletion or base tree arc contraction to create another LGT network *N*′|*T*′, we cannot create a new clusters in the base tree, i.e., every cluster of *T*′ is also in *T*. In fact, transfer deletions preserve all clusters, and a base tree arc contraction of (*u, v*) removes the cluster of *v* and preserves every other cluster.

Now let *v* be a bad vertex of *N*_1_|*T*_1_, and let *u* be the unique vertex such that [*u, v*] is a tree pair. Notice that any LGT reduction of 𝒩_1_ that contains the cluster 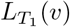 cannot be isomorphic to an LGT reduction of *N*_2_|*T*_2_. By our above observations, this means that *v* needs to be contracted eventually. Before this can be achieved, every transfer arc with an attachment point on [*u, v*] needs to be deleted. Counting these deletions, plus the contraction of (*u, v*), amounts to *w*(*v*) operations. This is true for every bad vertex, and one can easily see that any minimum sequence of operations can be altered, if necessary, so that it first deletes every bad transfer arc, then contracts every bad vertex with its parent. The same must be done with *N*_2_|*T*_2_. The number of operations needed to perform this is

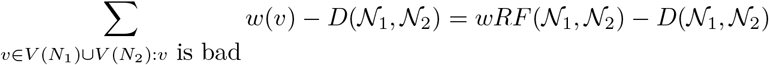

where we subtract *D*(𝒩_1_, 𝒩_2_) because in the summation, every doubly bad transfer arc is counted twice (and those that are bad but not doubly bad are counted once). Once this step is done, we obtain LGT reductions 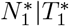 and 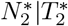 of *N*_1_|*T*_1_ and *N*_2_|*T*_2_, respectively, as described in our statement. Note that 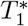 and 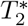 now have the same clusters. They are thus homeomorphic, and all that remains is to apply transfer deletions to make the LGT networks make isomorphic. By definition, this requires 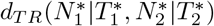 additional operations. This shows that any minimum sequence must perform at least the number of operations in our statement, and that number can easily be seen to also be an upper bound, since it corresponds to deleting bad transfers, contracting bad vertices, and handling remaining transfers.

#### Theorem 2

*The d*_*LGT*_ *distance satisfies all conditions of a metric*.

*Proof*. The fact that *d*_*LGT*_ (*N*_1_|*T*_1_, *N*_2_|*T*_2_) = 0 ⇔ *N*_1_|*T*_1_ ≃ _*LGT*_ *N*_2_|*T*_2_ holds by definition, since a common LGT reduction can be reached by 0 operations if and only if the two networks are already LGT-isomorphic. Symmetry is also easily verified.

Let us focus on the triangle inequality. We want to show that for any triple of LGT networks 𝒩_1_ = *N*_1_|*T*_1_, 𝒩_2_ = *N*_2_|*T*_2_, 𝒩_3_ = *N*_3_|*T*_3_ on the same leaves, we have

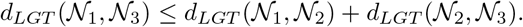

Let 𝒩_12_ = *N*_12_|*T*_12_ be a maximum common LGT reduction of 𝒩_1_, 𝒩_2_ and let 𝒩_13_ = *N*_13_|*T*_13_ be a maximum common LGT reduction of 𝒩_2_, 𝒩_3_. Observe that to reach a common LGT reduction of 𝒩_1_ and 𝒩_3_, we can first transform 𝒩_1_ into 𝒩_12_ and 𝒩_3_ into 𝒩_23_, and then find a common LGT reduction of the two LGT reductions. Therefore we get:

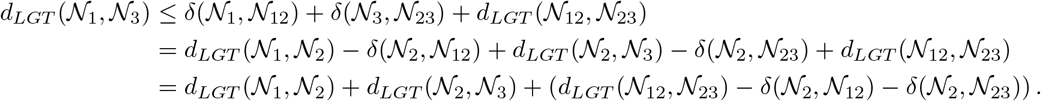

We next claim that *d*_*LGT*_ (𝒩_12_, 𝒩_23_) ≤ *δ*(𝒩_2_, 𝒩_12_) + *δ*(𝒩_2_, 𝒩_23_), which will prove the triangle inequality. To see this, we view 𝒩_12_ and 𝒩_23_ as obtained from 𝒩_2_ by applying operations, which lets us assume that *V* (*N*_12_) ⊆ *V* (*N*_2_) and *V* (*N*_23_) ⊆ *V* (*N*_2_) (this is without loss of generality, it just lets us avoid introducing an isomorphism in the notation). Also note that we may assume that to go from 𝒩_2_ to 𝒩_12_, all transfer deletions are done first, followed by all tree arc contractions (this is because if a contraction is followed by a transfer deletion, we can just move the transfer deletion before since contractions do not create nor destroy transfers). The same holds for turning 𝒩_2_ into 𝒩_12_. Let *A*_12_ (resp. *A*_23_) be the set of transfer arcs deleted from 𝒩_2_ to 𝒩_12_ (resp.𝒩_23_). Likewise, let *C*_12_ (resp. *C*_23_) be the set of arcs of the base tree contracted from 𝒩_2_ to 𝒩_12_ (resp. 𝒩_23_) after all transfer deletions were applied. Then, consider the LGT reduction 𝒩′ = *N*′|*T*′ obtained from 𝒩_2_ by deleting every transfer arc in *A*_12_ ∪ *A*_23_, and then contracting every base tree arc in *C*_12_ ∪ *C*_23_ (note that these arcs have no attachment points since we delete all transfer arcs first).

Notice that 𝒩′ is an LGT reduction of 𝒩_12_, since we can start from 𝒩_12_ and delete every arc from *A*_23_ that it still contains, and then contract the base tree arcs from *C*_23_ that remain (note, these are well-defined since we assume that *V* (*N*_12_), *V* (*N*_23_) ⊆ *V* (*N*_2_)). Likewise, 𝒩′ can be obtained from 𝒩_23_ by deleting the transfer arcs in *A*_12_ and contracting the tree arcs in *C*_12_. Thus, 𝒩′ is a common LGT reduction of 𝒩_12_ and 𝒩_23_. The distance *δ*(𝒩_12_, 𝒩′) is at most *δ*(𝒩_2_, 𝒩_23_) since we apply at most|*A*_23_| +|*C*_23_| operations. Likewise, *δ*(𝒩_23_, 𝒩′) is at most *δ*(𝒩_2_, 𝒩_12_). We thus get the desired claim, since

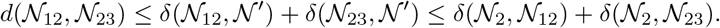

### B Proofs for Section 4 (The complexity of computing *d*_*T R*_ (and *d*_*LGT*_))

#### Theorem 3

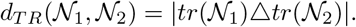

*Proof*. We first prove our identity on *d*_*TR*_(𝒩_1_, 𝒩_2_), and then argue the linear time complexity.

**Proof that** *d*_*TR*_(𝒩_1_, 𝒩_2_) = |*tr*(𝒩_1_)Δ*tr*(𝒩_2_)|. Consider the LGT network 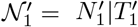 obtained from 𝒩_1_ by deleting every transfer arc that is not in 𝒩_2_. Formally, this means we delete every transfer arc (*u, v*) of 𝒩_1_ such that *u* is on [*u*_1_, *u*_2_], *v* is on [*v*_1_, *v*_2_] and ([*u*_1_, *u*_2_], [*v*_1_, *v*_2_]) ∈ *tr*(𝒩_1_) \ *tr*(𝒩_2_). Let 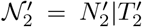 be obtained from 𝒩_2_ analogously, by deleting every transfer arc corresponding to an element of *tr*(𝒩_2_) *tr*(𝒩_1_). One can easily see that every such transfer must be deleted to obtain a common transfer reduction, because transfer deletions cannot create new transfers between tree pairs, so|*tr*(𝒩_1_)Δ*tr*(𝒩_2_)| is a lower bound on *d*_*LGT*_ (𝒩_1_, 𝒩_2_). We argue that it is also an upper bound.

We claim that 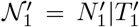 and 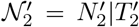 are LGT-isomorphic, which proves this upper bound. Let us first prove that 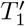 and 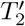 are isomorphic. Since *T*_1_ and *T*_2_ were assumed to be homeomorphic, and since transfer deletions can only remove attachment points in those trees, and not tree vertices, we know that 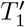 and 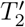 are still homeomorphic. Thus [*u, v*] is a tree pair of 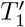 if and only if [*u, v*] is a tree pair of 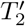. If the two trees are not isomorphic, it would thus mean that there is some tree pair [*u, v*] of 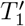 with one attachment point on it, but [*u, v*] in 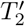 has no attachment point on it (or the roles of 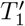 and 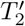 could be swapped, with the same arguments). This means that every transfer arc of 𝒩_1_ with an endpoint on [*u, v*] is in *tr*(𝒩_1_) *tr*(𝒩_2_), because if *tr*(𝒩_2_) had the same transfer arc, it would still be in 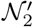 and the attachment point on [*u, v*] would have remained. There is therefore a contradiction, since every transfer arc of 𝒩_1_ with an attachment point on [*u, v*] should be deleted, and the attachment point should have been contracted. We therefore deduce that 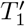 and 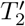 are isomorphic.

Let us assume for simplicity that 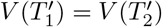, and that 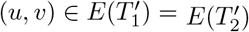 (we say for simplicity here, because the correspondence between vertices should technically rely on an isomorphism, which we omit). It remains to show that (*u, v*) is a transfer arc of 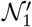 if and only if (*u, v*) is a transfer arc of 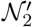. This follows from the fact that there is at most one attachment point between each tree pair. In more detail, if, for transfer arc (*u, v*) of 𝒩_1_, *u* is on [*u*_1_, *u*_2_] and *v* is on [*v*_1_, *v*_2_], then *u* must be the single attachment point on [*u*_1_, *u*_2_] and *v* the single attachment point on [*v*_1_, *v*_2_]. Because the pair ([*u*_1_, *u*_2_], [*v*_1_, *v*_2_]) is in both *tr*(𝒩_1_) and *tr*(𝒩_2_), there is also a transfer arc in 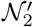 whose tail is on [*u*_1_, *u*_2_] and whose head is on [*v*_1_, *v*_2_]. There is only one option for the the tail and head of this transfer, which are *u* and *v*, the single attachment points between the two tree pairs. Thus (*u, v*) is also in 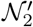. One can argue in the same manner that every transfer in 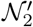 is in 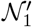, and therefore the two networks are LGT-isomorphic. This shows that *d*_*TR*_(𝒩_1_, 𝒩_2_) = |*tr*(𝒩_1_)Δ*tr*(𝒩_2_)|.

#### Proof of complexity

Now let us consider the time complexity of computing *d*_*LGT*_ (*N*_1_|*T*_1_, *N*_2_|*T*_2_). The reader may refer to the skeleton of the algorithm in Algorithm 1 and Algorithm 2, shown later in the appendix. We assume that all vertices of *N*_1_ and *N*_2_ are uniquely labeled with an integer between 1 and *n*_1_ + *n*_2_, where *n*_*i*_ = |*V* (*N*_*i*_)| for *i* ∈ {1, 2}. Note that *n*_1_ + *n*_2_ is in *O*(*m*_1_ + *m*_2_). We assume that we have access to both base trees *T*_1_ and *T*_2_, and that the list of transfer arcs of both networks are stored separately from the base trees.

The algorithm performs the following tasks:

1. Find the bad vertices (corresponding to bad clusters) and compute *wRF* (𝒩_1_, 𝒩_2_). To do this, we take the base trees *T*_1_, *T*_2_ and contract their attachment points, which can be done in time *O*(*n*_1_ + *n*_2_) (since attachment points are subdivision vertices, each contraction just removes two arcs and adds another). On the two resulting trees, we run Day’s algorithm, which in addition to computing the standard RF distance also finds the bad vertices in linear time *O*(*n*_1_ + *n*_2_) [39]. We can use a table with *n*_1_ + *n*_2_ bits, one entry for each possible vertex, with a 0 at position *i* if vertex *i* is not bad, and a 1 if it is bad. In this way, we can know in constant time whether a vertex is bad when traversing the base trees. We can then easily calculate the weight of bad vertices by just counting the number of transfers on the attachment point above them, and calculate the *wRF* component in linear time (the counts can be obtained by traversing the lists of transfers and incrementing a counter for their two respective ends).
2. Delete all the bad transfer arcs. For this we traverse the base trees and, whenever a bad vertex *u* is encountered, for each attachment point *v* just above *u*, we delete all transfer arcs that include *v* from the list of transfers. If we encounter a transfer arc that was already deleted, it is doubly bad and we can increment *D*(𝒩_1_,𝒩_2_). Doing all the necessary deletions and calculating *D*(𝒩_1_,𝒩_2_) in linear time requires a bit of care. We can proceed as follows. We take the list of bad vertices and sort it. Since vertex indices are at most *n*_1_ + *n*_2_, sorting takes linear time with bucket sort, for instance. We denote this list by *B*. Then, the lists of transfers of 𝒩_1_ and 𝒩_2_ store pairs of the form (*i, j*), where *i, j* are integers bounded by *n*_1_+*n*_2_. The two lists have at most *m*_1_+*m*_2_ elements. We can thus use radix sort to sort the two lists of transfers in time *O*(*m*_1_ + *m*_2_ + *n*_1_+*n*_2_), which is *O*(*m*_1_+*m*_2_) (using lexicographic order). We can then obtain the list of bad transfers *bad*_*tail*_ whose tail is in *B*, by maintaining an index in *B* and an index in the transfer list, and retaining each bad transfer whose first coordinate is a the current position in *B*, and advancing appropriate indices (an example of this strategy is used in Algorithm 4). Similarly, we can obtain the list of bad transfers *bad*_*head*_ whose head is in *B*, by sorting the transfer lists with respect to the second coordinate and repeating the process. Next, we compute the size of the intersection of *bad*_*tail*_ and *bad*_*head*_ to obtain *D*(𝒩_1_, 𝒩_2_) (to get this size, sort the two lists and do the two-index trick again). Finally, we can remove the bad transfers from our lists, again by sorting and, in a single pass, deleting every element that occurs in *bad*_*tail*_, then doing a second pass to delete those that are in *bad*_*head*_. Every step described here takes time *O*(*m*_1_ + *m*_2_). It is then not difficult to detect the now unused attachment points in both base trees and contract them.
3. Contract all bad vertices. If we do a pre-order traversal of the base trees and contract bad vertices when they are encountered, each vertex of either base trees changes parent at most once, so this takes time *O*(*n*_1_ + *n*_2_).
4. Compute *d*_*TR*_(𝒩_1_, 𝒩_2_) = |*tr*(𝒩_1_) Δ*tr*(𝒩_2_)| on the resulting networks. For this, we represent a transfer arc (*u, v*) as two tree pairs [*u*_1_, *u*_2_] and [*v*_1_, *v*_2_]. For each network and each transfer arc (*u, v*), we just look at the parent and child of *u* in the base tree to get [*u*_1_, *u*_2_], and do the same on *v* to get [*v*_1_, *v*_2_] (here, we use the assumption that each tree pair has at most one attachment point). We store *tr*(𝒩_1_) and *tr*(𝒩_2_) in two lists, which contain elements of the form ([*i, j*], [*k, l*]). These are quadruples of integers between 1 and *n*_1_ + *n*_2_, and again can be sorted in linear time with radix sort. Once *tr*(𝒩_1_) and *tr*(𝒩_2_) are sorted, we can maintain an index on each list to compute the size of their intersection in linear time, and thus get the size of the symmetric difference. This is detailed in Algorithm 4.

Since each step takes time *O*(*m*_1_ + *m*_2_) to calculate, we achieve the desired complexity.

##### Theorem 4

*Proof*. We prove the correctness of the reduction given above.

#### Time-consistency

We first verify that the networks are time-consistent. We build time maps *λ*_1_ for *N*_1_|*T*_1_ and *λ*_2_ to *N*_2_|*T*_2_. First note that we only add attachment points to arcs in *E*, which are arcs of the initial tree *T* whose heads are leaves. Thus, if we take the subgraph induced by the ancestors of the tails of arcs in *E* (inclusively) in either *N*_1_|*T*_1_ or *N*_2_|*T*_2_, we have a tree. It is easy to assign times in *λ*_1_ and *λ*_2_ such that all tails of arcs in *E* have time less than 0 (their ancestors will have negative times, which we allow for simplicity). It therefore suffices to assign a time above 0 to attachment points in a time-consistent manner.

In *N*_1_|*T*_1_, assign in *λ*_1_ a time of 4 to the two ends of every *t*_*x*_[*i*] transfer arc, and a time of 5 to the two ends of each *f*_*x*_[*i*] transfer arc. Now consider a clause *c* and recall that on arc *e*_*c*_, we added three attachment points *v*_*c*_[0], *v*_*c*_[1], *v*_*c*_[2]. Those correspond to the variables *x*_*c*,0_, *x*_*c*,1_, *x*_*c*,2_, which we recall are ordered so that positive literals appear first. Assign a time to the *v*_*c*_[*j*] vertices that correspond to positive literals a time in {1, 2, 3}, and a time to those for negative vertices a time of 6 or more (by our ordering, this can be done in a time-consistent manner). Finally, for each transfer arc whose head is a *v*_*c*_[*j*] vertex, assign the tail the same time as *v*_*c*_[*j*].

Let us argue that *λ*_1_ labeling is time-consistent. By construction, the ends of each transfer arc have equal times. Then, for *i* ∈ [0; *d*(*x*) − 1], there are exactly 2 attachment points on *e*_*x*_[*i*], namely *t*_*x*_[*i*].tail and *f*_*x*_[*i* − 1].head, with *t*_*x*_[*i*].tail above *f*_*x*_[*i*].head. Those have time-labels by *λ*_1_ equal to 4 and 5, respectively, which is time-consistent. On *e*_*x*_[*i*′] for *i* ∈ [0; *d*(*x*) − 1], there are exactly 3 attachment points: *t*_*x*_[*i*].head followed by *f*_*x*_[*i*].tail, again assigned respective times 4 and 5. There is a third attachment point *a*_*c,x*_, whose time is either in {1, 2, 3} or greater than 5, corresponding respectively to a positive occurrence of *x* in *c* (in which case *a*_*c,x*_ is above *t*_*x*_[*i*].head) and a negative occurrence of *x* in *c* (in which case *a*_*c,x*_ is below *f*_*x*_[*i*]). In both cases, the ordering of time labels is respected. Our construction of *λ*_1_ shows that for each tree pair corresponding to an arc of *E*, the times of the attachment points go in increasing order, and it follows that *N*_1_|*T*_1_ is time-consistent.

Now let us build time map *λ*_2_ for *N*_2_|*T*_2_. For each *i* ∈ [0, *d*(*x*) − 1], set the time of both ends of *f*_*x*_[*i*] to 1, and the time of both ends of *t*_*x*_[*i*] to 5. Then for each clause *c* and arc *e*_*c*_, the attachment points are, in order, *v*_*c*_[2], *v*_*c*_[1], *v*_*c*_[0], which we assign respective times 2, 3, 4. The tail of each transfer ending at one of these *v*_*c*_[*j*] vertices is assigned the same time as *v*_*c*_[*j*].

To see that this makes *N*_2_|*T*_2_ time-consistent, for *e*_*x*_[*i*] with *i* ∈ [0; *d*(*x*) − 1], the attachment points occuring on this arc have times 1 and 5. For 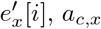 is below *f*_*x*_[*i*].tail and above *t*_*x*_[*i*].head, and is assigned one of the times in {2, 3, 4}. This is between 1 and 5, and so the times along 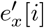 are in increasing order. This concludes the proof of time-consistency.

We next show that *ϕ* is satisfiable if and only if a common transfer reduction can be achieved using a total of 2*N* + 4*m* deletions.

#### *ϕ* satisfiable ⇒ existence of a large common reduction

Consider an assignment of each variable to true or false that satisfies *ϕ*. We show how to delete transfers from both networks, as illustrated in Figure 9. If a variable *x* is set to true, we remove the transfers *f*_*x*_[*i*] for all *i* ∈ [0; *d*(*x*) −1] in both networks. If it is set to false, likewise we remove *t*_*x*_[*i*] for *i* ∈ [0; *d*(*x*) − 1] in both networks. This amounts to 2 ∑_*x*_ *d*(*x*) = 2*N* deletions. Then, for each clause *c* of *ϕ*, we pick a variable *x* whose assignment satisfies *ϕ* (as true if *x* appears positively, as false otherwise) and remove in both networks all transfers arriving at *e*_*c*_, except the one originating at *a*_*c,x*_. For each clause, a total of 4 transfer deletion occurs this way in both networks (we delete two incoming transfers per clause per network). We have thus performed 2*N* + 4*m* deletions.

**Fig. 8:**
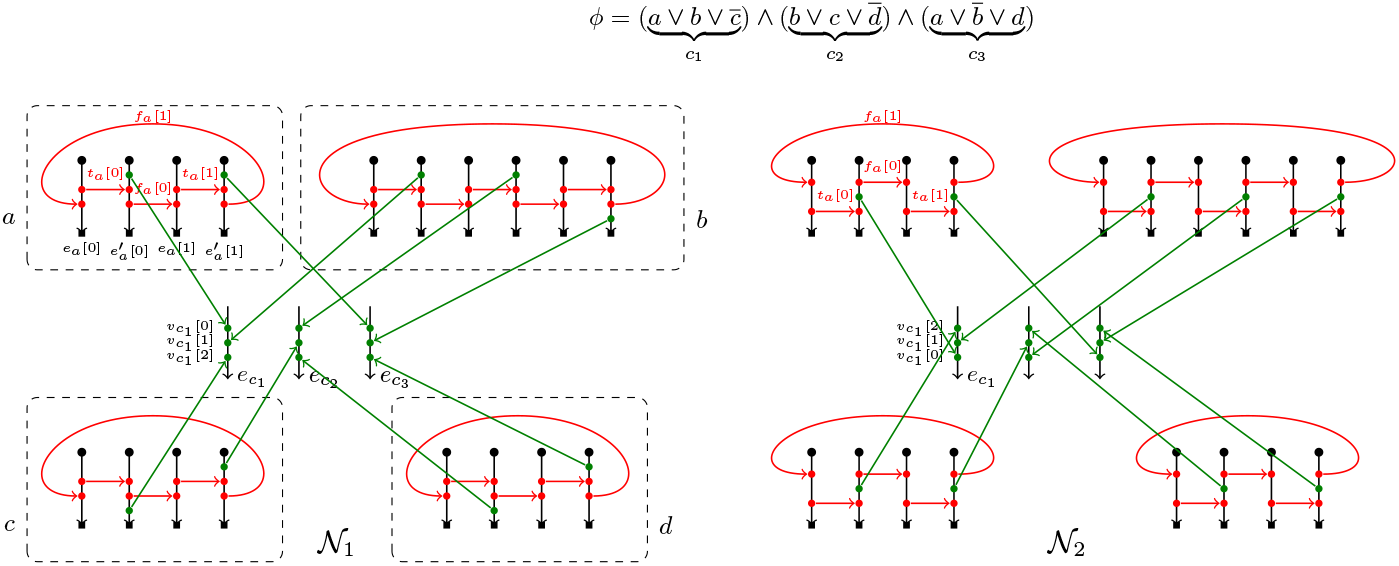
Example of the reduction (same figure as in the main text).

**Fig. 9:**
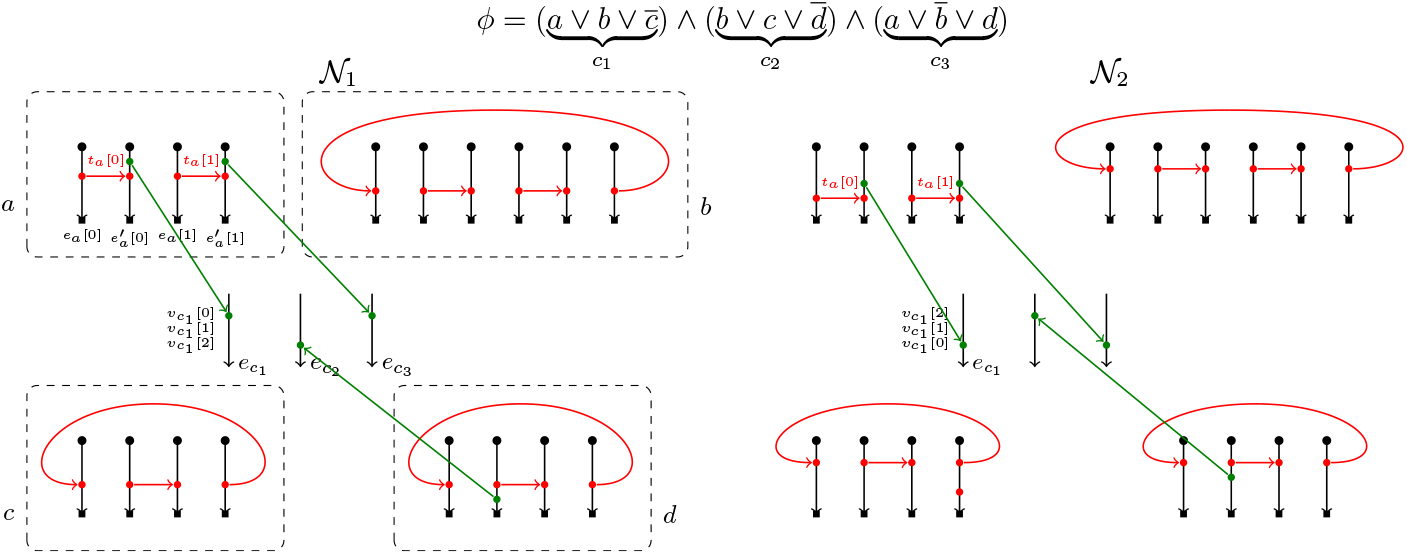
The solution obtained in the first direction, corresponding the the satisfying assignment that puts *a* = *true, b* = *c* = *d* = *false*. Here, *c*_1_ and *c*_3_ are satisfied by *a* = *true* and *c*_2_ by *d* = *false*.

We now argue that after these deletions on *N*_1_|*T*_1_ and *N*_2_|*T*_2_, we have achieved the same LGT network. Note that the two networks are identical above the tails of *E*, so we must only consider the transfer arcs between the tree pairs corresponding to arcs of *E*. To start with, for each variable *x*, either the set {*t*_*x*_[*i*]}_*i*∈ [0;*d*(*x*) − 1]_ or {*f*_*x*_[*i*]}_*i*∈[0;*d*(*x*) − 1]_ is kept, and the same choice is made in both networks, following the variable assignments. Therefore, between the tree pairs corresponding to the *e*_*x*_[*i*] and 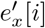 arcs, for all *x*, there is in the reductions of both networks exactly one transfer between *e*_*x*_[*i*] and 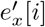 for *i* ∈ [0; *d*(*x*) − 1] in the former case, and between 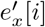 and *e*_*x*_[*i* + 1] in the latter (addition modulo *d*(*x*)). Moreover, if *x* is the variable chosen to satisfy *c*, and *c* is the *i*-th clause in which *x* appears, there is also *a*_*c,x*_ kept on 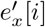. If *x* appears positively in *c*, then *a*_*c,x*_ is above *t*_*x*_[*i*].head on 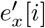 in *N*_1_. This is also the case in *N*_2_, regardless of whether *x* appears positively or negatively in *c*. Likewise, if *x* appears negatively in *c*, then *a*_*c,x*_ is below *f*_*x*_[*i*].tail in *N*_1_. This is also the case in *N*_2_, again regardless of whether *x* appears positively or negatively in *c*. As for the edges *e*_*c*_ for each clause *c*, exactly 1 attachment point remains on it in each network, and is the end-point of a transfer originating from the same arc *e*_*x*_[*i*], with *x* the variable that was picked to witness that *c* is satisfied, and with *c* the *i*-th clause in which *x* appears. Therefore, the reductions applied to *N*_1_ and *N*_2_ have given two networks in which transfers occur between exactly the same set of pairs of edges. In addition, the ordering of the attachment points of these transfers is also the same in both networks. Overall, a common reduction has been reached, with a total of 2*N* + 4*m* transfer deletions.

#### Existence of a large common reduction ⇒ *ϕ* Satisfiable

Conversely, suppose that a common transfer reduction *M* has been obtained from *N*_1_|*T*_1_ and *N*_2_|*T*_2_ with less than 2*N* + 4*m* transfer deletions. We need to show that *ϕ* is satisfiable. This direction is a bit more complicated and we introduce a useful tool before proceeding.

##### Transferring above other transfers

Let *N T* be an LGT network. We call a pair of attachment points *uv* a *transfer pair* if either (*u, v*) or (*v, u*) is a transfer arc of *N*. Note, the *uv* notation emphasizes that the direction is ignored. Given two tree pairs [*x, y*] and [*z, t*] of the base tree *T*, and *e* = [*p, q*] another tree pair of *T*, we say that [*x, y*] is *has a transfer above* [*z, t*] *in e* if there are 2 transfer pairs *ab, cd* such that (1) *a* is on [*x, y*] (2) *c* is on [*z, t*] (3) *b* and *d* are on *e* with *d* ≺_*T*_ *b*. We denote this relationship by [*x, y*] ⇒_*e*_ [*z, t*]. Note that we may have both [*x, y*] ⇒_*e*_ [*z, t*] and [*z, t*] ⇒_*e*_ [*x, y*] in a network with base tree *T*. This notion is illustrated on Figure 10.

**Fig. 10:**
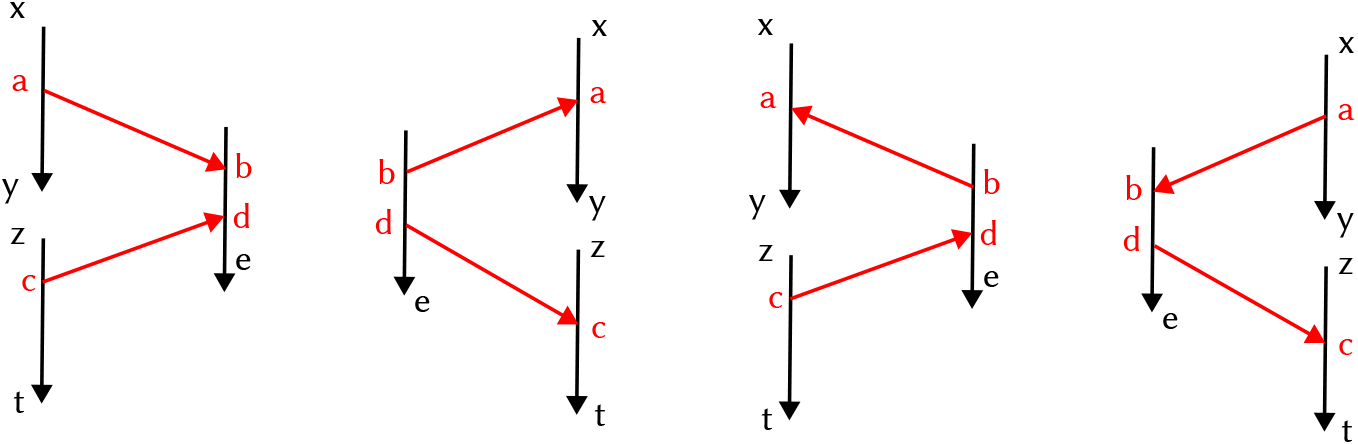
The different ways the relation [*x, y*] ⇒_*e*_ [*z, t*] can materialize. [*x, y*] and [*z, t*] essentially play the role of labels of the end points of the transfers. With this point of view, the relation [*x, y*] ⇒_*e*_ [*z, t*] simply states that “label [*x, y*] is above label [*z, t*] in *e*”.

When taking reductions of a network, the following property stating that ⇒_*e*_ relations cannot be created will be useful:

*Property 1*. Let 𝒩′ = *N*′|*T*′ be a transfer reduction of 𝒩 = *N*|*T*. If we do not have [*x, y*] ⇒_*e*_ [*z, t*] in *N*|*T*, then we also do not have [*x, y*] ⇒_*e*_ [*z, t*] in *N*′|*T*′.

*Proof*. We prove the result if 𝒩, 𝒩′ differ by 1 transfer deletion, and the result follows by induction. We show the contraposition of our statement, i.e., that given two tree pairs [*x, y*] and [*z, t*] such that [*x, y*] ⇒_*e*_ [*z, t*] for some tree pair *e* in 𝒩′, we also have [*x, y*] ⇒_*e*_ [*z, t*] in 𝒩. Let (*u, v*) be the transfer that is deleted to go from 𝒩 to 𝒩′. We also denote *s*_1_*t*_1_ and *s*_2_*t*_2_ transfer pairs in *N*′|*T* such that *s*_1_ is on [*x, y*], *t*_1_ is on *e, s*_2_ is on [*z, t*], *t*_2_ is on *e* and *t*_2_ ⪯_*T*_ *t*_1_ in 𝒩′ (such transfer pairs exist since [*x, y*] ⇒_*e*_ [*z, t*] in 𝒩′). To finish, we denote the transfer arc sets of 𝒩 and 𝒩′ by *E*_*t*_ and 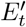, respectively. We get that 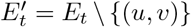, and 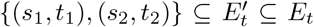. We also have *t*_2_ ⪯_*T*_ *′ t*_1_ in 𝒩. Indeed, recall that to go from 𝒩′ to 𝒩, two arcs of *E*(*T*′) are possibly subdivided to create *u* and *v*, and then an arc is added between *u* and *v*. Whether one of the subdivided arcs is on the path from *t*_1_ to *t*_2_ in *T*′ or not, we have *t*_1_ ⪯_*T*_ *t*_2_ in 𝒩 as well. Overall, [*x, y*] ⇒_*e*_ [*z, t*] in 𝒩 as well.

Let us now go back to our main proof. Property 1 will be used to show that if [*x, y*] ⇒_*e*_ [*z, t*] holds in 𝒩_1_ but not in 𝒩_2_, then we must apply a deletion that makes [*x, y*] ⇒_*e*_ [*z, t*] become false in 𝒩_1_ (since the property says that the ⇒_*e*_ relation can never be created in 𝒩_2_).

We first argue that any common transfer reduction of 𝒩_1_ and 𝒩_2_ requires *at least* 2*N* + 4*m* deletions. Let us first note that a transfer arc *t*_*x*_[*i*] is deleted in #x1D4A9;_1_ if and only if that same transfer arc *t*_*x*_[*i*] is deleted in 𝒩_2_, since this is the only transfer arc between the tree pairs of *e*_*x*_[*i*] and 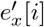. The same holds for the *f*_*x*_[*i*] arcs.

Next, note that for any variable *x* and for *i* ∈ [0; *d*(*x*) 1], 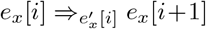 in 𝒩_1_ (as *t*_*x*_[*i*].head is above *f*_*x*_[*i*].tail on 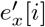 in 𝒩_1_), and not the reverse, whereas 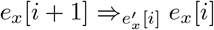 in 𝒩_2_ (*t*_*x*_[*i*].head is below *f*_*x*_[*i*].tail on 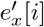 in 𝒩_2_), and not the reverse (as usual, additions are to be taken modulo *d*(*x*)). In both cases, the reverse relation is not true, so if both *t*_*x*_[*i*] and *f*_*x*_[*i*] are kept in a common transfer reduction 𝒩 = *N*|*T* of 𝒩_1_ and 𝒩_2_, we would have both 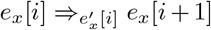 and 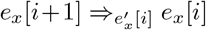 in 𝒩, contradicting Property 1. Likewise, for *i* ∈ [0; *d*(*x*) − 1], if both *f*_*x*_[*i*] and *t*_*x*_[*i* + 1] are kept in a common transfer reduction 𝒩, then we have both 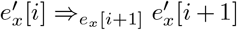 and 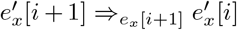 in 𝒩, contradicting Property 1.

In other words, any two transfers that have their attachment point on the same tree pair are incompatible: one of the two must be deleted. This yields at least 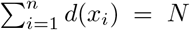 deletions in the variable gadgets in each network. In addition, we show that if exactly *N* deletions occur in a variable gadget associated to *x*, then either {*t*_*x*_[*i*]}_*i* ∈ [0;*d*(*x*) −1]_ or {*f*_*x*_[*i*]}_*i*∈ [0;*d*(*x*) − 1]_ is entirely kept. If exactly *N* deletions occur, then for each *x* and for each *i* ∈ [0; *d*(*x*) − 1], either *t*_*x*_[*i*] or *f*_*x*_[*i*] is removed but not both. Likewise, either *f*_*x*_[*i*] or *t*_*x*_[*i* + 1] must be deleted, but not both. Suppose *t*_*x*_[0] is kept, then *f*_*x*_[*d*(*x*) − 1] and *f*_*x*_[0] are removed. This implies *t*_*x*_[1] is kept. Iterating this argument shows that exactly {*t*_*x*_[*i*]} _*i*∈[0;*d*(*x*) −1]_ is kept. We can show similarly that if only *N* deletions occur and *f*_x_ [0] is kept in the clause gadget of *x*, then {*f*_x_ [*i*]}_*i*∈[0;*d*(*x*)−1]_ is kept.

As for the clause gadget, consider a clause *c*, and two variables *x* = *x*_*c,j*_ and 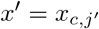 appearing in *c*. We suppose w.l.o.g. that *x* is before *x*′, that is *j < j*′. We denote *i* and *i*′ the integers such that *c* is the *i*-th clause in which *x* appears, and the *i*′-th clause in which *x*′ appears (so, *c* = *c*_*x,i*_ = *c*_*x*_*′*_,*i*_*′*). Note that we have 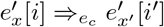 in 𝒩_1_ and not the reverse, while the opposite is true in 𝒩_2_, i.e., 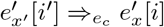 in 𝒩_2_ because we reversed the order of attachment points on *e*_*c*_. As this is true for any two variables *x, x*′ that appear in *c*, it implies through Property 1 that at most 1 transfer entering *e*_*c*_ may be kept in each network. It implies at least 2*m* deletions in each network. In addition, if only 2*m* deletions occur, exactly one transfer arriving in each *e*_*c*_ must be kept.

Finally, let *x* be a variable in which we kept all *t*_*x*_[*i*] arcs. One can see that if, in the gadget for *x*, we kept an arc of 𝒩_1_ from some *a*_*c,x*_ vertex to some *v*_*c*_[*j*] vertex, then *x* must occur positively in *c*. This is because if the occurrence was negative, *a*_*c,x*_ would be below *t*_*x*_[*i*].head in 𝒩_1_ but above *t*_*x*_[*i*].head in 𝒩_2_ and we would again violate Property 1. For the same reasons, if all *f*_*x*_[*i*] arcs were kept instead, then all remaining arcs starting from *a*_*c,x*_ vertices correspond to negative occurrences. Thus in a common transfer reduction, all clause arcs *e*_*c*_ whose incoming transfer arcs are from the same *x* gadget have the same type of occurrence, all positive or all negative. This means that we can assign *x* = *true* if the *t*_*x*_[*i*]’s were kept and *x* = *false* if the *f*_*x*_[*i*]’s were kept. Since every *e*_*c*_ arc has an incoming transfer from some *x* gadget that corresponds to satisfying *c*, this assignment shows that *ϕ* is satisfiable.

#### Proof that computing *d*_*T B*_ is NP-hard

##### Construction of the proof of hardness for *d*_*TB*_

We now modify the reduction described in Section 4.2 to show that *d*_*TB*_ is NP-hard. Letting *ϕ* be an instance of 3-SAT, we let 𝒩_1_ = *N*_1_|*T*_1_ and 𝒩_2_ = *N*_2_|*T*_2_ be the LGT networks constructed from *ϕ* as described previously. Recall that both have a base tree that originated from a tree *T* that contains a set of arcs *E*, all having their head vertex being a leaf, to which we added attachment points.

Let us obtain a tree-based network 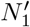 from *N*_1_. Let (*u, v*) be an arc in *E*, and note that [*u, v*] is a tree-pair of *N*_1_|*T*_1_. Now make the transformation to *N*_1_ that is illustrated in Figure 11. That is, denoting by *p*_*u*_ the parent of *u* in *N*_1_ (and *T*_1_), the arc (*p*_*u*_, *u*) is replaced with a subgraph with 5 vertices where *u* is now a reticulation (in the figure, the possible *u* vertices are the larger black circles — *p*_*u*_ is not shown, but instead of having *u* as a child it would have the root of the inserted subnetwork in blue as a child). Denote by 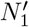 the network obtained from *N*_1_ after applying this transformation to every (*u, v*) in *E*. Likewise, denote by 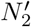 the network obtained from *N*_2_ by applying the same transformations. It is not difficult to show that 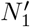 and 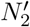 are tree-based networks. Moreover, the point of this transformation is that the set of transfer arcs of our original reduction are enforced.

**Fig. 11:**
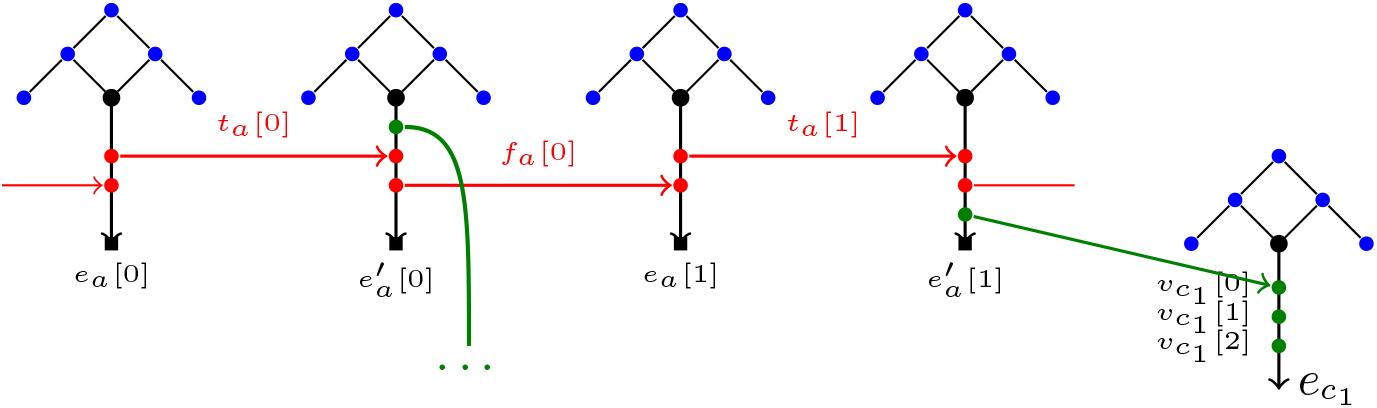
Modification of the gadgets of *N*_1|_*T*_1_ for a variable *a* and a clause *c*_1_. For each tail *u* of an arc in *E* with parent *p*_*u*_ (*p*_*u*_ is not shown): we remove the arc (*p*_*u*_, *u*), we add the subgraph consisting of the five blue vertices as shown, then add the root of that subgraph as a child of *p*_*u*_. In each such subgraph, the two dangling vertices are newly introduced leaves.

**Fig. 12:**
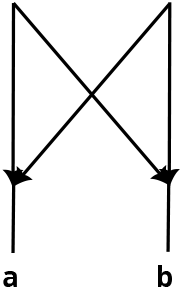
The pattern shown the left is not allowed for the sub-class of orchard networks (see for instance [26] for a definition). Although it can be considered non-realistic (it is not time-consistent). Yet, the network prediction methods used in Sections 5.2 and 5.3 (namely [34] and [31]) could in principle return networks exhibiting this pattern, requiring comparison methods that are capable of comparing ideally all tree-based networks [36], and not just orchard networks.

**Fig. 13:**
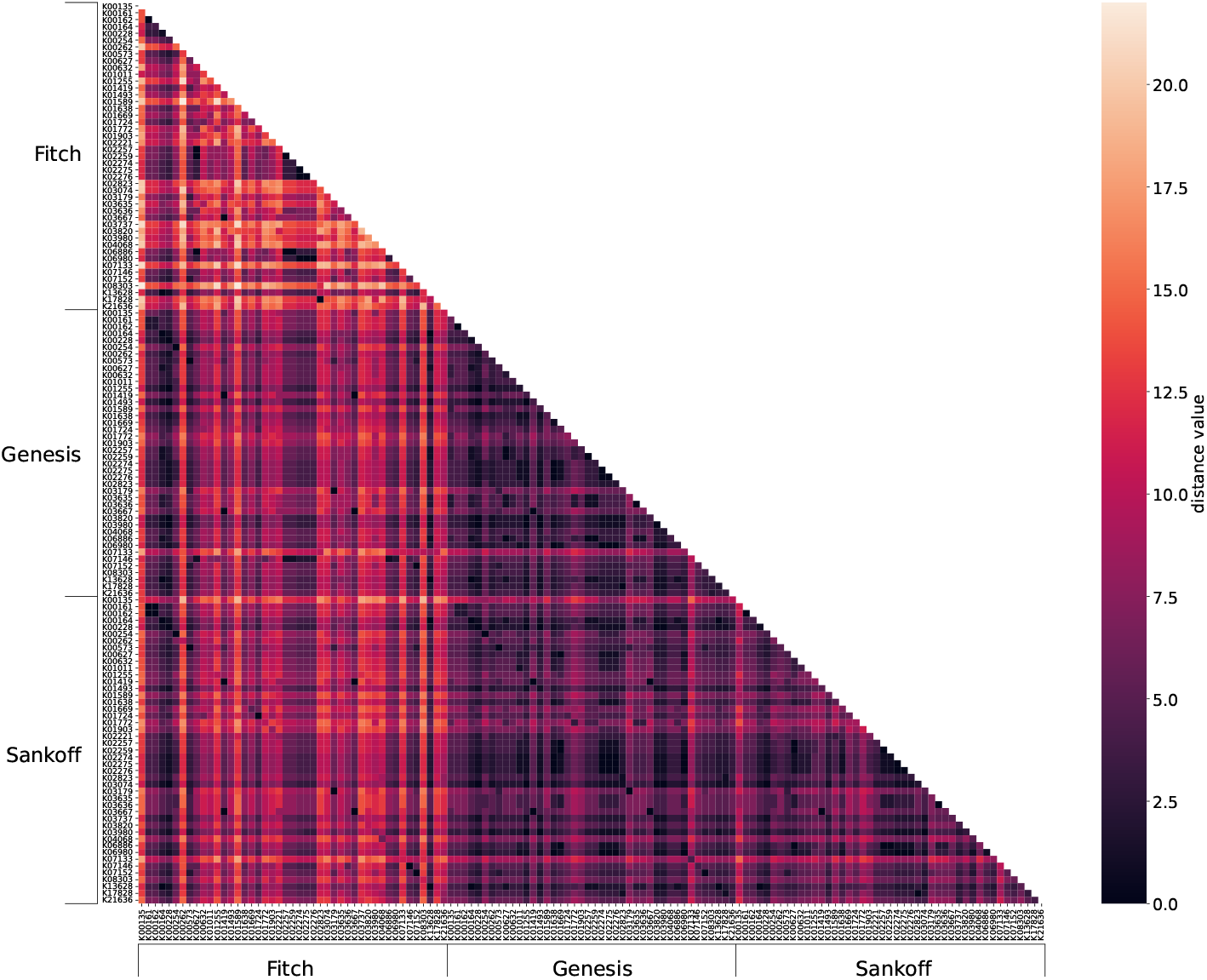
Ancestral character reconstruction obtained using the Basic, Sankoff, and Genesis algorithms.

###### Lemma 1

*Let* (*x, y*) *be a transfer arc of N*_1_|*T*_1_. *Then for any base tree* 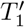 *in* 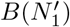, *the arc* (*x, y*) *is a transfer arc of* 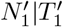. *Likewise, if* (*x, y*) *is a transfer arc of N*_2_|*T*_2_, *then for any base tree* 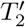 *in* 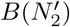, *the arc* (*x, y*) *is a transfer arc of* 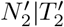.

*Proof*. We prove the result for *N*_1_|*T*_1_ first. Let 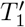 be a base tree of 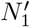. Let *e*_*c*_ ∈ *E* be an arc of the initial tree corresponding to a clause *c*, and let *e*_*c*_ = (*u, v*). Note that [*u, v*] is a tree pair of *N*_1_|*T*_1_ on which we added attachment points *v*_*j*_[0], *v*_*j*_[1], *v*_*j*_[2]. Because in 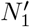, *u* is a reticulation, the arc (*u, v*_*j*_[0]) must be in the base tree 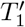 of 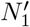 (recall that by definition, the base tree must have the same leaves as 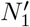, and so every non-leaf of 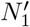 needs to have at least one child in the base tree). Thus, the other arc entering *v*_*c*_[0], namely some (*a*_*c,x*_, *v*_*c*_[0]) arc, cannot also be in the base tree and thus it must be a transfer of 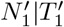. Following the same logic, (*v*_*c*_[0], *v*_*c*_[1]) must be in 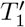 and the other arc entering *v*_*c*_[1] is a transfer arc, and the same holds for *v*_*c*_[2]. Thus, the transfer arcs with an end on *e*_c_ are the same in *N T* and 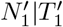.

We next argue that each *t*_*x*_[*i*] arc cannot be in the base tree. Consider the arc 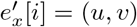 and note that *t*_*x*_[*i*].head is on the tree pair corresponding to 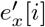. By construction, in *N*_1_ either *t*_*x*_[*i*].head has *u* as a parent, or has an attachment point *a*_*c,x*_ for a clause *c* as a parent. In the former case, (*u, t*_*x*_[*i*].head) must be in the base tree because *u* has one child, implying that *t*_*x*_[*i*] cannot also be in the base tree. In the latter case, the arc from *a*_*c,x*_ to *e*_*c*_ was argued to be a transfer, so (*a*_*c,x*_, *t*_*x*_[*i*].head) must be in the base tree since *a*_*c,x*_ needs a child, implying in turn that *t*_*x*_[*i*] cannot be in the base tree. In all cases, *t*_*x*_[*i*] must be a transfer arc. Now consider a *f*_*x*_[*i*] arc. In *N*_1_, *f*_*x*_[*i*].head has *t*_*x*_[*i*+ 1].tail as a parent. Since we now know that *t*_*x*_[*i*] is not in the base tree, then (*t*_*x*_[*i*].tail, *f*_*x*_[*i*].head) is in the base tree and *f*_*x*_[*i*] must be a transfer arc. Thus, all transfer arcs of *N*_1_|*T*_1_ are also transfers of 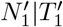.

For *N*_2_|*T*_2_, we use similar ideas to argue that any base tree of *N*_2_ results in the same transfer arcs. The transfer arcs entering *e*_*c*_ cannot be in any base tree of *N*_2_ for the same reasons as before. Each *f*_*x*_[*i*] cannot be in the base tree because each reticulation *u* on the upper end of *e*_*x*_[*i* + 1] must have *f*_*x*_[*i*].head as its child. Finally, each *t*_*x*_[*i*] cannot be in the base tree, because in 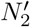 the parent of *t*_*x*_[*i*].head is some *a*_*c,x*_ attachment point, which has its arc going into *e*_*c*_ a transfer and which therefore needs *t*_*x*_[*i*].head as its child, preventing *t*_*x*_[*i*] from being in the base tree.

###### Theorem 5

*Computing d*_*TB*_ *is NP-hard*.

*Proof*. We propose a reduction from 3-SAT. Given an instance *ϕ* of 3-SAT, first construct the networks 𝒩_1_ = *N*_1_|*T*_1_ and 𝒩_2_ = *N*_2_|*T*_2_ from Section 4.2, and then apply the modifications described above to obtain tree-based networks 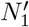 and 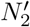, resulting in the *d*_*TB*_ instance. We claim that 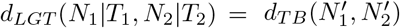, which is sufficient to prove NP-hardness since it implies *ϕ* is satisfiable if and only if 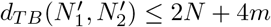 as before.

Let us first show that 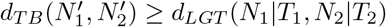. Let 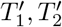 be base trees of *N*_1_, *N*_2_ such that 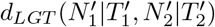 is minimum. Let *D*_1_ be the set of transfer arcs deleted from 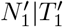 and let *D*_2_ be those deleted from 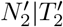 to obtain LGT networks 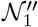 and 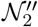 that form a maximum common LGT reduction (note, contractions could have been performed, but we don’t even count them). Recall that by Lemma 1, all transfer arcs in *N*_1_|*T*_1_ and *N*_2_|*T*_2_ are also in 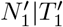 and 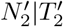, respectively. Consider the LGT networks obtained from *N*_1_|*T*_1_ by deleting all of its transfers that are also in *D*_1_, and the LGT network obtained from *N*_2_|*T*_2_ by deleting all those that are also in *D*_2_, resulting in new LGT networks 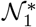 and 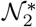. As a result, the subgraph induced by the descendants of tails of elements from *E* is identical in 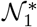 and in 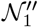, and identical in 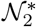 and in 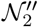 (recall that 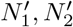 are obtained by replacing the arcs incoming into the tails of elements from *E* by a subgraph, so everything “below” tails of *E* was unaltered, and thus we get the same subgraphs in these parts after making the same deletions). Moreover, since 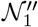 and 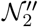 are LGT-isomorphic, the subgraphs induced by the descendants of tails of elements from *E* are identical in both LGT networks, and thus the same holds for 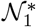 and 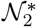. It follows that 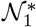 and 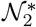 are themselves LGT-isomorphic. Since we made at most 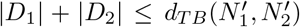 transfer deletions to reach 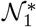 and 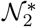, we get 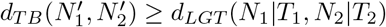.

We show the converse inequality 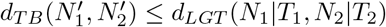. In 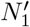, we take the base tree 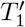 with the same transfer arcs as 𝒩_1_, and for each reticulation above *E* we choose the transfer arc arbitrarily. For 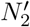, we take the base tree 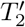 with the same transfer arcs as 𝒩_2_, and for the reticulations above *E* we make the same arbitrary choice as in 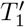. It then becomes easy to see that a sequence of transfer deletions that makes 𝒩_1_ and 𝒩_2_ LGT-isomorphic can also be used to make 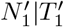 and 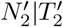 LGT-isomorphic. This yields 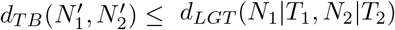.

#### Proof that *d*_*LGT*_ is FPT in the number of transfers

We recall our proposition of interest here.

##### Proposition 1

*Proof*. Using the ideas in the proof of Theorem 3, one may calculate *wRF* and *D*(_1, 2_) in linear time, and it remains to state how to calculate *d*_*TR*_(𝒩_1_,𝒩_2_). Suppose that 𝒩_1_ has *t*_1_ transfer arcs and 𝒩_2_ has *t*_2_. One may iterate over all 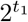 subsets of transfers of 𝒩_1_ and all 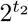 sets of transfers of 𝒩_2_ in time 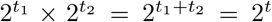. For each subset, we can obtain transfer reductions 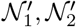 in linear time (technically, we must copy each network before doing so, which just adds another linear time term to the complexity). Then, one needs to check whether the two reductions are LGT-isomorphic. This may also be done in linear time as follows. We already know at this point that the two networks are homeomorphic, so we only need to check that the transfer arcs are identical. For that, one may simply store, for each transfer arc (*u, v*): (1) the tree pairs [*u*_1_, *u*_2_] and [*v*_1_, *v*_2_] that *u* and *v* are on; (2) the rank of *u* on [*u*_1_, *u*_2_] and the rank of *v* on [*v*_1_, *v*_2_] (i.e., the number of attachment points to traverse on the tree pair to reach the vertex). We then ensure LGT-isomorphic by checking, for each transfer arc (*u, v*) in either network, that the corresponding transfer arc with the same extra information exists in the other network. To do this, assuming that vertices are labeled 1 to *n*_1_ + *n*_2_, with *n*_*i*_ = |*V* (*N*_*i*_)| as in the proof of Theorem 3, we can store each transfer arc in the form ([*i, j*], [*k, l*], *r, r*′), where *r, r*′ are the aforementioned ranks. This is a sextuple with integers bounded by *n*_1_ + *n*_2_. These can be sorted in linear time using radix sort, and then we can check that the lists of transfers are identical.

##### Theorem 6

*Given two binary LGT networks* 𝒩_1_,𝒩_2_ *both of level at most ℓ, one can compute d*_*TR*_(𝒩_1_,𝒩_2_) *and thus d*_*LGT*_ (𝒩_1_,𝒩_2_) *in time O*(4^*ℓ*^ · *m*^2^), *where m is the total number of edges in both networks*.

#### Proofs for Theorem 6 (FPT in the level)

If the input networks are binary, we can do better and obtain an FPT algorithm in the *level* of the input networks [11]. To define this notion, first, an arc (*u, v*) of an LGT network *N*|*T* is a *bridge* if deleting (*u, v*) from *N* makes its underlying undirected graph disconnected. We call *v* a bridge-end. A *blob* of *N* is a maximal subgraph that contains no bridge. The *level* of 𝒩 (and *N*) is the maximum number of reticulations in a blob of *N*. In binary networks, the level can be much lower than the number of transfers, making it a useful parameter in practice.

We say that a pair of LGT networks 𝒩_1_ = *N*_1_|*T*_1, 2_ = *N*_2_|*T*_2_. forms a *clean instance* if the transfer cleaning rule is not applicable on them. We can obtain a clean instance by applying the transfer cleaning rule until exhaustion.

Note that a transfer arc in either network cannot be a bridge, since there is always an undirected cycle that contains a transfer arc. For the same reason, a bridge-end cannot be an attachment point. Therefore, bridge-ends are tree vertices and have a single parent. This will be useful in the lemmas that build up to Theorem 6.

##### Lemma 2

*Suppose that* 𝒩_1_ = *N*_1_|*T*_1_, 𝒩_2_ = *N*_2_|*T*_2_ *is a clean instance. Then v is a bridge-end of N*_1_ *if and only if v is a bridge-end of N*_2_.

*Proof*. Let (*u, v*) be a bridge of *N*_1_. As mentioned just above, *v* is a tree vertex of 𝒩_1_ and thus *v* is also in 𝒩_2_. If *v* is also a bridge-end of *N*_2_, we are done, so assume otherwise for contradiction. Thus in the undirected version of *N*_2_, the root can reach some vertex *w* descending from *v*, say through a path *P*. Without loss of generality, we may assume that *w* is a tree vertex, since otherwise we redefine *w* to be its next descendant that is a tree vertex. Thus *w* is also a vertex of *N*_1_.

We can partition *P* so that it is equal to *P*_1_ − *Q*_1_ − *P*_2_ − *Q*_2_ … − *Q*_*h*−1_ − *P*_*h*_ where: (1) each *P*_*i*_ is a subpath of *T*_2_ that starts and ends at a tree vertex (so it uses no transfer arc); (2) each *Q*_*i*_ starts with the last vertex *p*_*i*_ of *P*_*i*_, goes to an attachment point on an edge incident to *p*_*i*_, takes a transfer arc (in either direction), and goes to one of the tree vertices from the tree-pair that the transfer is on. We argue that in the undirected version of *N*_1_, the root can also reach *w* without going through *v*. To see this, observe that for each *P*_*i*_, the undirected version of *T*_1_ also contains a path 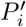 with the same ends that avoid *v* (we just traverse the same tree pairs as *P*_*i*_, so *P*_*i*_ and 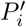 only differ by the attachment points they go through). Likewise, for each *Q*_*i*_, the undirected version of *N*_1_ also has a path 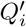 with the same ends as *Q*_*i*_, that uses a transfer, and whose interior vertices are all attachment points (the reason is that the instance is clean, so the tree pairs having the transfer arc used by *Q*_*i*_ also have a transfer in *N*_1_). Moreover, 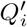 also avoids *v*. It follows that the sequence of edges formed by 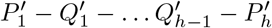 goes from the root of *N*_1_ to *w* without going through *v*. That sequence may not be a path because it could repeat vertices, but in this case we can remove the cycles in that sequence to obtain a path. We have reached a contradiction on (*u, v*) being a bridge of *N*_1_, so the bridge-ends of *N*_1_ are also bridge-ends of *N*_2_. The converse holds using the same arguments.

Recall that a blob of a network *N* is a maximal subgraph *B* of *N* that contains no bridge. A *pendant bridge of B* is a bridge (*u, v*) such that exactly one of *u* or *v* is in *V* (*B*) (if *v* is not in it, then the bridge goes “out”, and if *u* is not in it, then *v* is the root of the blob and the bridge is entering the blob). The set of pendant bridges of *B* is denoted *pend*(*B*).

##### Lemma 3

*Suppose that* 𝒩_1_ = *N*_1_|*T*_1_, 𝒩_2_ = *N*_2_|*T*_2_ *is a clean instance. Let B*_1_ *(resp. B*_2_*) be the blob of N*_1_ *(resp. N*_2_*) that contains the root. Then v is the end of a pendant bridge of B*_1_ *if and only if v is the end of a pendant bridge end of B*_2_.

*Proof*. Let (*u, v*) be a pending bridge of *B*_1_, with *v* the pendant bridge-end, and recall that *v* is a tree vertex. Thus *v* is also in *N*_2_. Suppose that *B*_2_ does not have a pending bridge with end *v*. By Lemma 2, *v* is also a bridge-end of *N*_2_, and so the arc (*u*′, *v*) of *N*_2_ whose head is *v* is a pending bridge of some blob 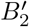 of *N*_2_ that is different from the root blob *B*_2_. Let (*p, q*) be the arc entering this blob 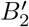 (this arc exists and is unique, see [49]). Then (*p, q*) is a bridge of *N*_2_ and therefore *q* is a bridge-end of *N*_1_, say on the bridge (*p*′, *q*) by Lemma 2. In *N*_2_, (*p, q*) separates the root from *v* and its parent, and because *v* descends from *q* in *N*_1_ as well, in *N*_1_ we have that (*p*′, *q*) separates the root from *v* and its parent *u*. This is a contradiction, since in *N*_1_ the vertex *u* should not be in the same blob as the root. This concludes the proof.

We are now setup to describe a dynamic programming algorithm that processes each blob independently. To this end, for an LGT network 𝒩 = *N*|*T* and a bridge-end *u*, let (*u*) 𝒩 = *N* (*u*)|*T* (*u*) denote the subnetwork of 𝒩 rooted at *u*, where *N* (*u*) (resp. *T* (*u*)) is the subgraph of *N* (resp. of *T*) induced by *u* and all its descendants. Note, by the definition of a bridge, no transfer arc of has one end in *N* (*u*) and the other outside.

Similarly, if *B* is a blob of *N* rooted at some vertex *u*, let *N B* denote the subgraph induced by ⟨*B*⟩ and all the vertices that are bottom ends of a pendant bridge of *B*. Note that these ends are the leaves of *N* ⟨*B*⟩.

##### Lemma 4

*Let* 𝒩_1_ = *N*_1_|*T*_1_, 𝒩_2_ = *N*_2_|*T*_2_ *be* binary *LGT networks that form a clean instance, and whose blobs containing the root are respectively B*_1_ *and B*_2_. *Then*

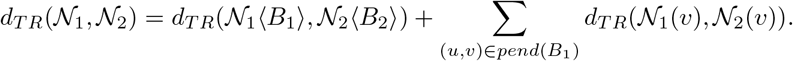

*Consequently, d*_*TR*_ *and thus d*_*LGT*_ *can be computed in time O*(4^*ℓ*^*m*^2^), *where ℓ is the maximum level of* 𝒩_1_ *or* 𝒩_2_ *and m* = |*E*(*N*_1_)| +|*E*(*N*_2_)|.

*Proof*. Let us first focus on the correctness of our claimed recurrences. By Lemma 3, the networks 𝒩_1_ ⟨*B*_1_⟩ and 𝒩_2_ ⟨*B*_2_⟩ have the same leaves. Hence, if we make those subnetworks isomorphic by transfer deletions, and then replace each of their leaves by isomorphic subnetworks, the result is an isomorphic network. Thus the recurrence gives a possible way to obtain a common transfer reduction, which justifies

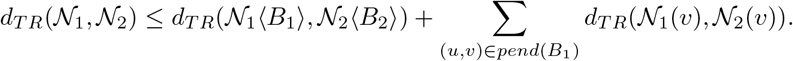

For the converse bound, let *R*_1_, *R*_2_ be a set of transfer arcs to remove from 𝒩_1_ and𝒩_2_, respectively, to make them isomorphic. These deletion sets must, in particular, make 𝒩_1_ ⟨*B*_1_⟩, 𝒩_2_ ⟨*B*_2_⟩ isomorphic, and for each end *v* of a pendant bridge, make 𝒩_1_(*v*) and 𝒩_2_(*v*) isomorphic. The aforementioned subnetworks share no transfer arc, so the subsets of *R*_1_, *R*_2_ within any such subnetwork are disjoint. This justifies

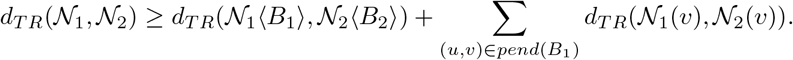

As for the complexity, we first make the networks clean by applying the transfer cleaning rule until exhaustion. This is easy to do in linear time *O*(*m*), using the techniques from Theorem 3. For concreteness, if we assume constant time operations on hash tables, we can find the transfers to delete in the preprocessing by hashing each transfer according to the tree pairs it is on, and then computing the symmetric difference in linear time. Once this is done, we can compute the *d*_*TR*_ entries by dynamic programming in a bottom-up fashion. It is easily seen that the only entries *d*_*TR*_(𝒩_1_(*v*), 𝒩_2_(*v*)) that will ever be required are those when *v* is the root of a blob (this includes the case *d*_*TR*_(𝒩_1_, 𝒩_2_), where *v* is taken to be the root of both networks). There are *O*(*n*) = *O*(*m*) such possible entries. For one specific entry *d*_*TR*_(𝒩_1_(*v*), 𝒩_2_(*v*)), we try deleting every subset of transfer arcs from the root blob of the first network, and every subset in the second. Since every reticulation is incident to at most one transfer arc in a binary LGT network, the number of transfer arcs is at most *ℓ* and there are thus 2^*ℓ*^ subsets to enumerate in each blob, for a total of 4^*ℓ*^ combinations. For each combination, we check whether applying the deletions leads to an isomorphic network, which adds linear time (see the proof of Proposition 1).

In summary, for each pair of blobs we spend time *O*(4^*ℓ*^*m*) to enumerate the deletion sets and test isomorphism on each, and there are *O*(*m*) entries to compute. The time for cleaning the networks does not impact this, so overall the time is *O*(4^*ℓ*^*m*^2^).

The complexity claimed in Theorem 6 then follows immediately from the previous lemma.

#### Algorithms for Section 4

Here we describe the algorithm from Proposition 1. The same ideas can be used for Theorem 6. The algorithm computes *d*_*LGT*_ as the sum of the weight of bad vertices (*wRF*), the number of extra transfer deletions to make (*D*_*tr*_), and the number of doubly bad transfers (*D*). Bad vertices are found using Day algorithm. Algorithm 1 integrates these components and returns the combined contribution of tree disagreement and transfer disagreement.

##### Algorithm 1

getD_LGT(*N*_1_|*T*_1_, *N*_2_|*T*_2_

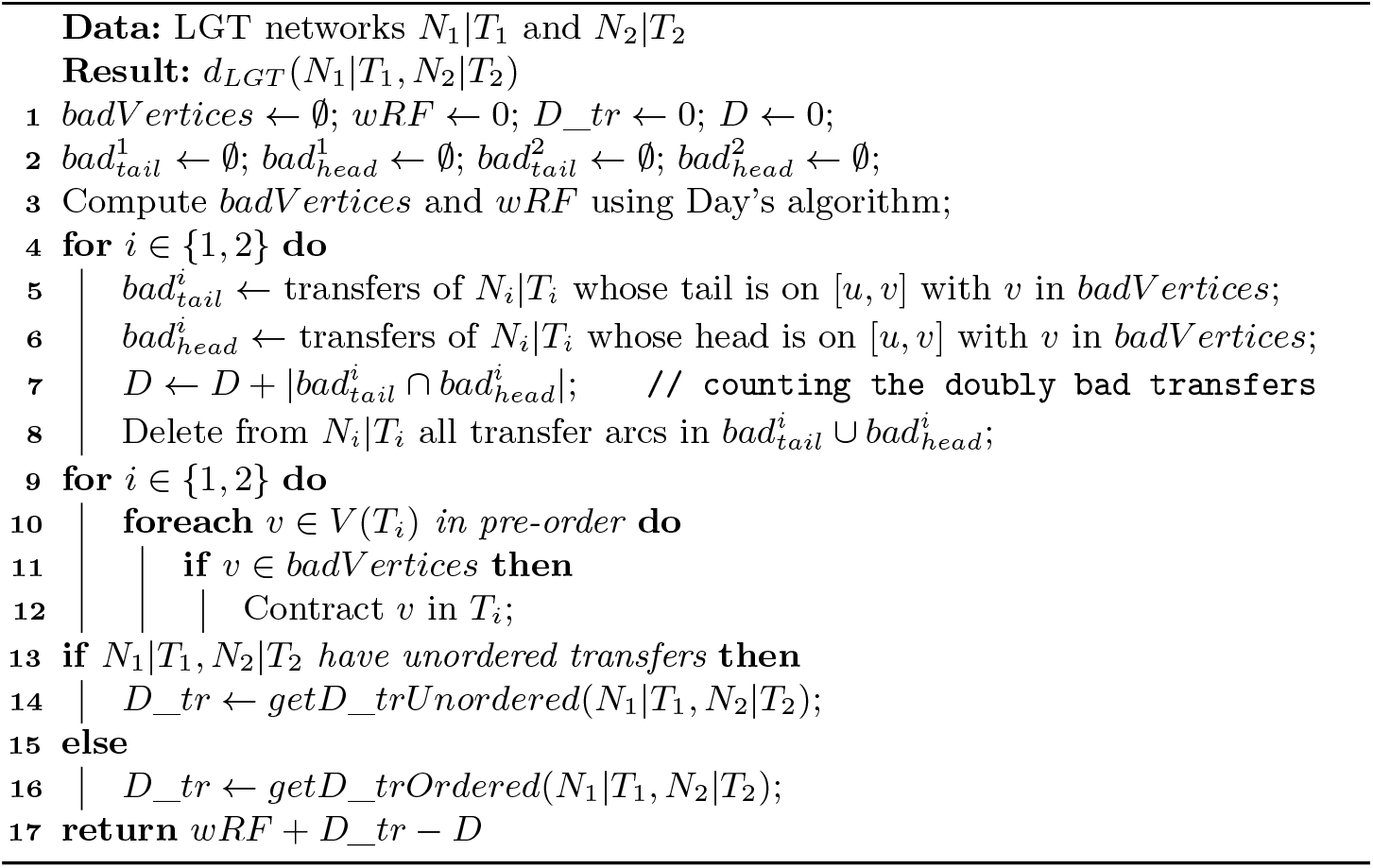

When transfer arcs are treated as unordered, the distance reduces to counting the arcs that appear in exactly one of the two networks. In this setting, no additional structural constraints need to be verified once the relative complements are computed. Algorithm 2 therefore returns the transfer distance directly as the total number of non-shared arcs. Note, the relative complement of *A* in *B* is the set difference *B\A*. We can obtain the symmetric difference *A* Δ *B* by computing both relative complements *A\B* and *B\A*. The algorithm for GetRelativeComplements is shown aferwards.

##### Algorithm 2

getD_trUnordered(*N*_1_|*T*_1_, *N*_2_|*T*_2_

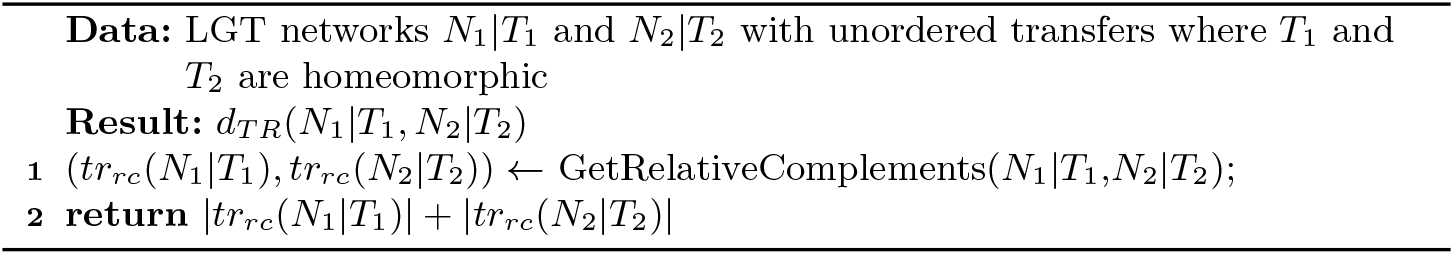

When transfers are ordered, our FPT algorithm simply explores all subsets of the remaining transfer arcs and finds the smallest pair of deletion sets that yields equivalent LGT structures, as shown in Algorithm 3.

##### Algorithm 3

getD_trOrdered(*N*_1_|*T*_1_, *N*_2_|*T*_2_)

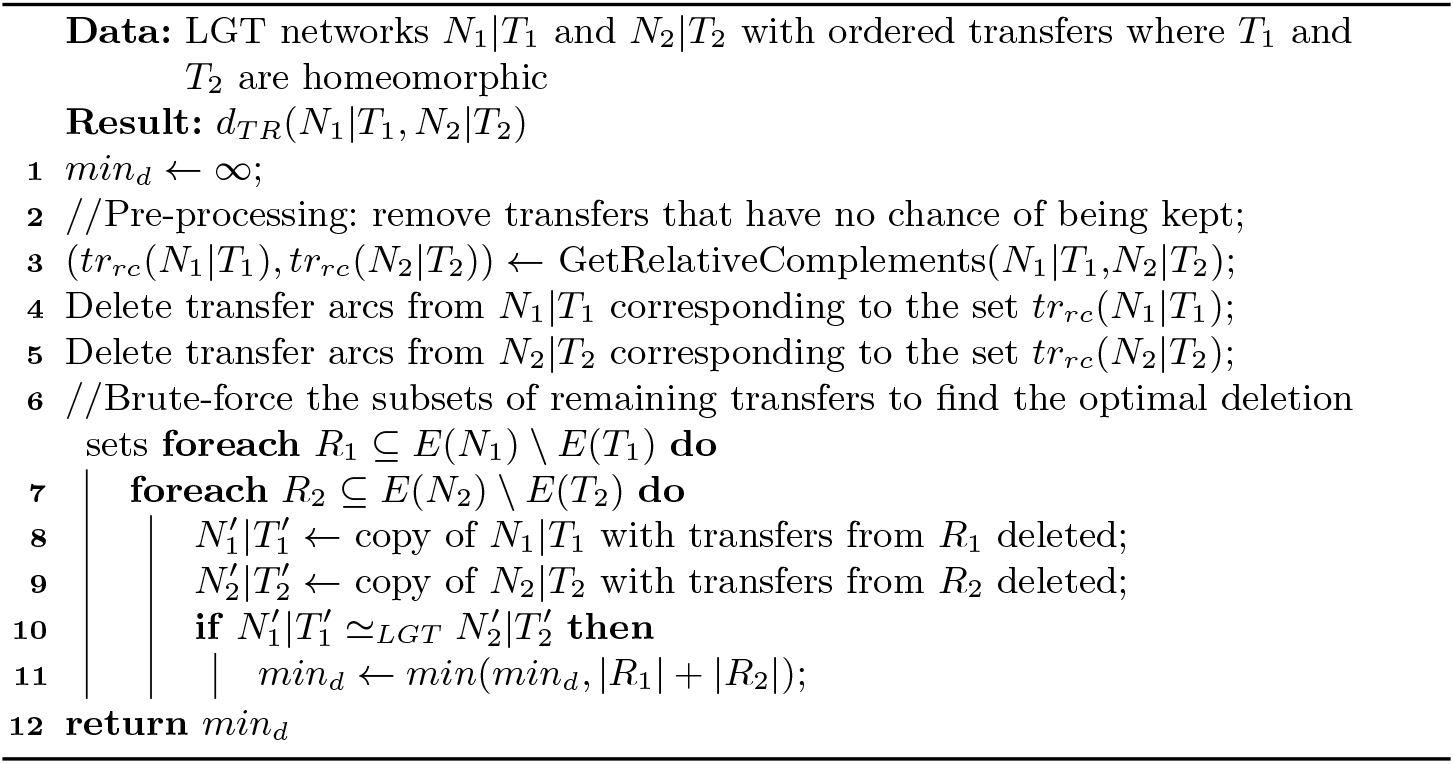

Algorithm 4 performs this computation of the relative complements, so that the returned pair of sets can be used to get the symmetric difference, or to delete the transfers in both networks. This is done by sorting both arc sets and scanning them in parallel to isolate the non-shared arcs. The transfers in *tr*(*N*_*i*_|*T*_*i*_) have the form ([*u*_1_, *u*_2_], [*v*_1_, *v*_2_]), which can be seen as a quadruple of integers between 0 and the sizes of the networks, in which case radix sort takes linear time.

##### Algorithm 4

getRelativeComplements(*N*_1_|*T*_1_, *N*_2_|*T*_2_)

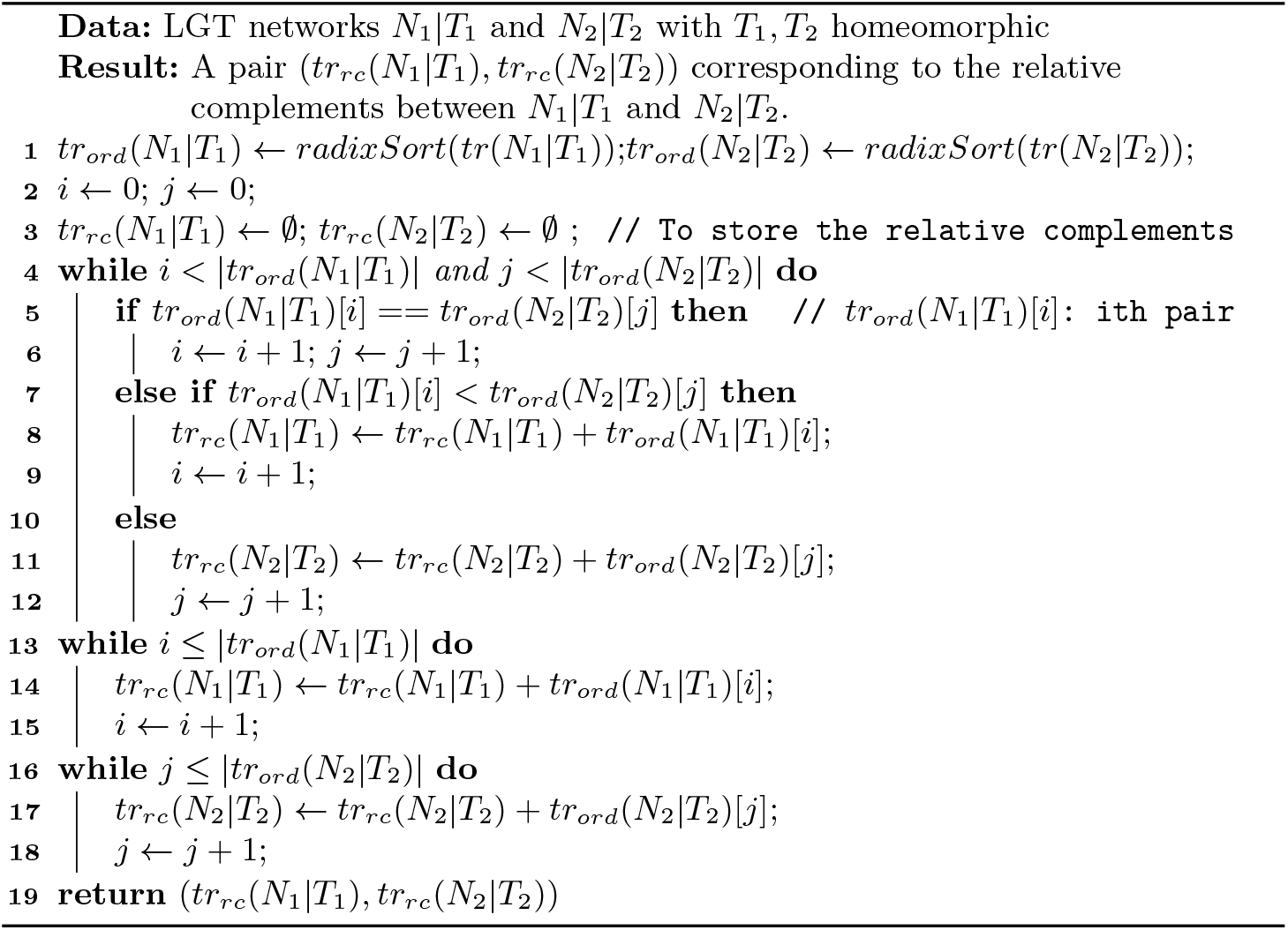

### C Proofs for Section 5 (Experiments)

